# Cell shape and antibiotic resistance is maintained by the activity of multiple FtsW and RodA enzymes in *Listeria monocytogenes*

**DOI:** 10.1101/589911

**Authors:** Jeanine Rismondo, Sven Halbedel, Angelika Gründling

**Author notes:** To whom correspondence should be addressed: Angelika Gründling –.

## Abstract

Rod-shaped bacteria have two modes of peptidoglycan synthesis: lateral synthesis and synthesis at the cell division site. These two processes are controlled by two macromolecular protein complexes, the elongasome and divisome. Recently, it has been shown that the *Bacillus subtilis* RodA protein, which forms part of the elongasome, has peptidoglycan glycosyltransferase activity. The cell division specific RodA homolog FtsW fulfils a similar role at the divisome. The human pathogen *Listeria monocytogenes* encodes up to six FtsW/RodA homologs, however their functions have not yet been investigated. Analysis of deletion and depletion strains led to the identification of the essential cell division-specific FtsW protein, FtsW1. Interestingly, *L. monocytogenes* encodes a second FtsW protein, FtsW2, which can compensate for the lack of FtsW1, when expressed from an inducible promoter. *L. monocytogenes* also possesses three RodA homologs, RodA1, RodA2 and RodA3 and their combined absence is lethal. Cells of a *rodA1/rodA3* double mutant are shorter and have increased antibiotic and lysozyme sensitivity, probably due to a weakened cell wall. Results from promoter activity assays revealed that expression of *rodA3* and *ftsW2* is induced in the presence of antibiotics targeting penicillin binding proteins. Consistent with this, a *rodA3* mutant was more susceptible to the β-lactam antibiotic cefuroxime. Interestingly, overexpression of RodA3 also led to increased cefuroxime sensitivity. Our study highlights that *L. monocytogenes* encodes a multitude of functional FtsW and RodA enzymes to produce its rigid cell wall and that their expression needs to be tightly regulated to maintain growth, cell division and antibiotic resistance.

**Importance:** The human pathogen *Listeria monocytogenes* is usually treated with high doses of β-lactam antibiotics, often combined with gentamicin. However, these antibiotics only act bacteriostatically on *L. monocytogenes* and the immune system is needed to clear the infection. Therefore, individuals with a compromised immune system are at risk to develop a severe form of *Listeria* infection, which can be fatal in up to 30% of cases. The development of new strategies to treat *Listeria* infections is therefore necessary. Here we show that the expression of some of the FtsW and RodA enzymes of *L. monocytogenes* is induced by the presence of β-lactam antibiotics and their combined absence makes bacteria more susceptible to this class of antibiotics. The development of antimicrobials that inhibit the activity or production of FtsW/RodA enzymes might therefore help to improve the treatment of *Listeria* infections and thereby lead to a reduction in mortality.

## Introduction

Bacterial cells are surrounded by a mesh of peptidoglycan (PG) that determines their shape and also protects the cells from lysis due to their high internal turgor pressure (Weidel and Pelzer, 1964, Vollmer et al., 2008, de Pedro and Cava, 2015). Peptidoglycan is comprised of glycan strands that are crosslinked by short peptides (Rogers et al., 1980). The glycan strands are composed of alternating N-acetylglucosamine and N-acetylmuramic acid residues that are connected by a β-1,4 glycosidic bond (Ghuysen and Strominger, 1963). The synthesis of peptidoglycan begins in the cytoplasm with the production of the PG precursor lipid II by the proteins MurABCDEF, MraY and MurG (Blumberg and Strominger, 1974, van Heijenoort, 2001, Scheffers and Pinho, 2005, Pinho et al., 2013). Lipid II is then transported across the cytoplasmic membrane by the flippase MurJ and Amj (Meeske et al., 2015, Sham et al., 2014, Ruiz, 2008) and subsequently incorporated in the growing glycan strand by glycosyltransferases. The polymerization and the crosslinking of the glycan strands are facilitated by the activity of glycosyltransferases and transpeptidases, respectively. Class A penicillin binding proteins (PBPs) are bifunctional enzymes that possess glycosyltransferase and transpeptidase activity, whereas class B PBPs only contain a transpeptidase domain (Höltje, 1998, Sauvage et al., 2008, Goffin and Ghuysen, 1998). In addition, some species such as *Escherichia coli, Staphylococcus aureus* and *Streptococcus pneumoniae* encode monofunctional glycosyltransferases (MGTs) that can also incorporate lipid II into the growing glycan strand (Park and Matsuhashi, 1984, Park et al., 1985, Karinou et al., 2018, Wang et al., 2001, Hara and Suzuki, 1984). *B. subtilis* encodes four class A PBPs and no MGT, however, deletion of all class A PBPs only manifests in small PG changes (McPherson and Popham, 2003). Recently, it has been shown that members of the SEDS (shape, elongation, division, sporulation) family of proteins, namely RodA and FtsW, also act as glycosyltransferases (Meeske et al., 2016, Cho et al., 2016, Emami et al., 2017, Taguchi et al., 2019). Both, RodA and FtsW, form complexes with cognate class B PBPs to enable polymerization and crosslinking of glycan strands (Cho et al., 2016, Taguchi et al., 2019, Leclercq et al., 2017, Fraipont et al., 2011, Reichmann et al., 2019). Interestingly, SEDS proteins and the class B PBPs are more conserved among different bacterial species than class A PBPs (Meeske et al., 2016).

In rod-shaped bacteria, peptidoglycan is synthesized by two multiprotein complexes, the elongasome that is essential for the cell elongation and the divisome that is crucial for the formation of the division septum (Nanninga, 1991, Carballido-Lopez and Formstone, 2007, Typas et al., 2012, Errington and Wu, 2017). RodA is part of the elongasome and is essential in many bacteria including *B. subtilis* and *S. pneumoniae* (Liu et al., 2017, Henriques et al., 1998). Depletion of RodA results in the production of enlarged, spherical cells in *B. subtilis* (Henriques et al., 1998). In contrast, FtsW is essential for cell division and cells depleted for FtsW grow as long filaments (Kobayashi et al., 2003, Boyle et al., 1997, Gamba et al., 2016). *B. subtilis* harbors a sporulation specific member of the SEDS family, SpoVE in addition to RodA and FtsW. SpoVE is dispensable for growth, however, it is essential for the synthesis of the spore cortex peptidoglycan (Henriques et al., 1992, Ikeda et al., 1989). Other *Bacillus* species such as *B. cereus* and *B. anthracis* possess 4 to 5 FtsW/RodA proteins and strains of different serotypes of the human pathogen *Listeria monocytogenes* encode even up to 6 FtsW/RodA homologs in their genome. However, their functions have not yet been investigated.

Here, we determined the role of the different FtsW and RodA homologs for the growth and cell morphology of *L. monocytogenes*. Our results show that *L. monocytogenes* encodes two FtsW and three RodA enzymes. Absence of either FtsW1 or of all three RodA proteins is lethal under standard laboratory conditions. *L. monocytogenes* infections are usually treated with high doses of β-lactam antibiotics such as ampicillin, which inhibit the transpeptidase activity of PBPs (Swaminathan and Gerner-Smidt, 2007). We demonstrate that the expression of two SEDS proteins, FtsW2 and RodA3, is induced in the presence of β-lactam antibiotics likely to compensate for the inhibition of PBPs and that a *rodA3* mutant is more sensitive to the β-lactam antibiotic cefuroxime. Antimicrobials inhibiting the activity of proteins of the SEDS family could therefore potentially improve the treatment of *Listeria* infections in the future.

## Materials and Methods

### Bacterial strains and growth conditions

All strains and plasmids used in this study are listed in Table S1. Strain and plasmid constructions are described in the supplemental materials and method section and all primers used in this study are listed in Table S2. *E. coli* strains were grown in Luria-Bertani (LB) medium and *L. monocytogenes* strains in brain heart infusion (BHI) medium at 37°C unless otherwise stated. If necessary, antibiotics and supplements were added to the medium at the following concentrations: for *E. coli* cultures, ampicillin (Amp) at 100 μg/ml and kanamycin (Kan) at 30 μg/ml, and for *L. monocytogenes* cultures, chloramphenicol (Cam) at 10 μg/ml, kanamycin (Kan) at 30 μg/ml and isopropyl β-D-1-thiogalactopyranoside (IPTG) at 1 mM. We used the *L. monocytogenes* strain 10403S and derivatives thereof. However, we refer to *L. monocytogenes* EGD-e gene and locus tag numbers as this was the first fully sequenced *L. monocytogenes* strain.

### Growth curves

Overnight cultures of wildtype *L. monocytogenes* 10403S and the indicated deletion strains were diluted to an OD_600_ of 0.01 or 0.05 in 15 ml BHI medium and the cultures were incubated at 37°C with shaking. Growth was monitored by determining OD_600_ readings at hourly intervals. For growth curves with the IPTG-inducible depletion strains 10403S Δ*ftsW1* i*ftsW* (ANG4314), 10403SΔ*ftsW1* i*ftsW2* (ANG5119) and 10403SΔ*rodA1-3* i*rodA1* (ANG5192), the strains were cultivated overnight in the presence of 1 mM IPTG. The next day, cells were washed once with fresh medium, diluted 1:50 in 5 ml BHI medium and grown for 8-10 h in the absence of the inducer. The cultures were diluted 1:100 into fresh BHI medium and grown until the next morning at 37°C. The depleted cells were then diluted to an OD_600_ of 0.01 and grown in the presence or absence of 1 mM IPTG at 37°C. Averages and standard deviations from three independent experiments were plotted.

### RNA extraction and quantitative real-time PCR (qRT-PCR)

For the extraction of RNA from *rodA* complementation strains, overnight cultures of *L. monocytogenes* strains 10403S, 10403SΔ*rodA1*Δ*rodA3*, 10403SΔ*rodA1*Δ*rodA3* pIMK3-*rodA1*, 10403SΔ*rodA1*Δ*rodA3* pIMK3-*rodA2* and 10403S Δ*rodA1*Δ*rodA3* pIMK3-*rodA3* were diluted in BHI medium (with 1 mM IPTG for the plasmid-containing complementation strains) to an OD_600_ of 0.05 and incubated at 37°C until the cultures reached an OD_600_ of 1. For the extraction of RNA from strain 10403SΔ*rodA1-3* i*rodA1* (ANG5192), bacteria were grown as described for the growth curve assay to deplete RodA1. Next, 10403S and depleted cells of 10403SΔ*rodA1-3* i*rodA1* were diluted to an OD_600_ of 0.01 and grown in BHI medium in the presence or absence of IPTG until an OD_600_ of 0.5. Twenty mls of the cultures were mixed with 47 ml guanidine thiocyanate (GTC) buffer (5 M GTC, 0.5% N-lauryl sarcosine, 0.1 M β-mercaptoethanol, 0.5% Tween-80, 10 mM Tris-HCl pH 7.5), bacteria were harvested by centrifugation and subsequently lysed using the FastRNA™ Pro Blue kit (MP Biomedicals). The total RNA was isolated by chloroform extraction and ethanol precipitation and further purified using the RNeasy mini kit (Qiagen) and finally treated with TURBO DNase (Invitrogen). cDNA was synthesized from 10 ng of RNA using the Superscript III first strand synthesis kit (Invitrogen). The expression of *rodA1, rodA2* and *rodA3* in the different strains was assessed using the TaqMan^®^ probe-based gene expression assay (Applied Biosystems). Expression of *gyrB* was used as control. The cycle threshold (Ct) values obtained for *rodA1, rodA2* and *rodA3* were normalized to the values obtained for *gyrB*. The fold changes of gene expression for the different strains were calculated using the ΔΔCt method.

### Determination of minimal inhibitory concentration (MIC)

The minimal inhibitory concentration for bacitracin, penicillin and moenomycin and lysozyme was determined using a microbroth dilution assay in 96-well plates. Approximately 10^4^ *L. monocytogenes* cells were used to inoculate 200 μl BHI containing two-fold dilutions of the different antimicrobials. The starting antibiotic concentrations were: 1 mg/ml for bacitracin A, 1 μg/ml for penicillin G, 0.8 or 1.6 μg/ml for moenomycin, 8 μg/ml cefuroxime and 10 mg/ml for lysozyme. The OD_600_ readings were determined after incubating the 96-well plates for 24 h at 37°C with shaking at 500 rpm in a plate incubator (Thermostar, BMG Labtech). The MIC value refers to the antibiotic concentration at which bacterial growth was inhibited by >90%.

### Determination of antibiotic susceptibility using a spot plating assay

Overnight cultures of the indicated *L. monocytogenes* strains were adjusted to an OD_600_ of 1 and 5 μl of serial dilutions were spotted on BHI agar plates or BHI agar plates containing 1 μg/ml cefuroxime and where indicated also 1 mM IPTG. Plates were photographed after incubation at 37°C for 24 h.

### Fluorescence and phase contrast microscopy

For bacterial cell length measurements, 100 μl of mid-log cultures were mixed with 5 μl of 100 μg/ml nile red solution to stain the cell membrane. Following incubation at 37°C for 20 min, the cells were washed once with PBS and resuspended in 50 μl PBS. 1.5 μl of the different samples were spotted on microscope slides covered with a thin agarose film (1.5 % agarose in distilled water), air-dried and covered with a cover slip. Phase contrast and fluorescence images were taken using a Zeiss Axio Imager.A1 microscope coupled to a AxioCam MRm and a 100x objective and processed using the Zen 2012 software (blue edition). For the detection of nile red fluorescence signals, the Zeiss filter set 00 was used. For the cell length determinations, 300 cells were measured for each experiment and the median cell length was calculated. Averages and standard deviations of the median cell length of three independent experiments were plotted.

### Peptidoglycan isolation and analysis

Overnight cultures of *L. monocytogenes* 10403S, 10403SΔ*rodA1*Δ*rodA3* and 10403SΔ*rodA1*Δ*rodA3* pIMK3-*rodA1* were used to inoculate 1 L BHI broth (with 1 mM IPTG for the complementation strain 10403SΔ*rodA1*Δ*rodA3* pIMK3-*rodA1*) to a starting OD_600_ of 0.06. The cultures were grown at 37°C until they reached an OD_600_ of 1, at which point the cultures were cooled on ice for 1 h. The bacteria were subsequently collected by centrifugation and peptidoglycan was purified and digested with mutanolysin as described previously (de Jonge et al., 1992, Corrigan et al., 2011). Digested muropeptides were analyzed by high-performance liquid chromatography (HPLC) and recorded at an absorption of 205 nm as described previously (de Jonge et al., 1992). For quantification, the areas of the main muropeptide peaks were integrated using the Agilent Technology ChemStation software. The sum of the peak areas was set to 100% and individual peak areas were determined. Averages and standard deviations from three independent extractions were calculated.

### β-galactosidase assay

For the determination of the β-galactosidase activity, overnight cultures of strains 10403S pPL3e-P*_lmo2689_-lacZ*, 10403SΔ*rodA1* pPL3e-P*_lmo2689_-lacZ* and 10403SΔ*rodA1*Δ*rodA2* pPL3e-*Pl_mo2689_-lacZ* were diluted 1:100 in fresh BHI medium and grown for 6 h at 37°C. Sample collection and preparation were performed as described previously (Gründling et al., 2004). Briefly, OD_600_ readings were determined (for the final β-galactosidase unit calculations) for the different cultures after 6 h of growth and cells from 1 ml culture were pelleted by centrifugation for 10 min at 13,200 × g, resuspended in 100 μl ABT buffer (60 mM K_2_HPO_4_, 40 mM KH_2_PO_4_, 100 mM NaCl, 0.1% Triton X-100, pH 7.0), snap frozen in liquid nitrogen and stored at −80°C until use. For the identification of substances inducing the expression of the *lmo2689-lmo2686* operon, an overnight culture of strain 10403S pPL3e-P*_lmo2689_-lacZ* was diluted 1:100 in fresh BHI medium and the culture incubated with shaking at 37°C until an OD_600_ of 0.5-0.6. The culture was divided into several flasks and incubated for two hours at 37°C in the presence or absence of the following substances: 0.5 μg/ml ampicillin, 0.05 μg/ml penicillin, 0.5 μg/ml vancomycin, 4 μg/ml cefuroxime, 0.05 μg/ml moenomycin, 0.5 mg/ml lysozyme, 1% ethanol, 300 μg/ml MgSO_4_ or 300 μg/ml EDTA. Bacteria were pelleted and samples frozen as described above.

Samples were thawed and 1:10 dilutions were prepared in ABT buffer. Fifty μl of the 1:10 diluted samples were mixed with 10 μl of 0.4 mg/ml 4-methyl-umbelliferyl-β-D-galactopyranoside (MUG) substrate prepared in DMSO and incubated for 60 min at room temperature (RT). A reaction with ABT buffer alone was used as negative control. Following this incubation step, 20 μl of each reaction was diluted into 180 μl of ABT buffer in a black 96-well plate and fluorescence values were measured using an HIDEX Sense Microplate Reader at 355 nm excitation and 460 nm emission wavelengths. 0.125-20 μM of the fluorescent 4-methylumbelliferone (MU) standard were used to obtain a standard curve. β-galactosidase units were calculated as (pmol of substrate hydrolyzed x dilution factor)/(ml culture volume x OD_600_ x minute). The amount of hydrolyzed substrate was determined from the standard curve as (emission reading – y intercept)/slope.

### Bacterial two-hybrid assays

Protein-protein interactions between the different FtsW/RodA homologs and class B PBPs were analyzed using the bacterial adenylate cyclase two-hybrid (BACTH) assay (Karimova et al., 1998). The indicated pUT18/pUT18c and pKT25 derivatives were co-transformed into *E. coli* strain BTH101. Transformants were selected on LB agar plates containing 100 μg/ml ampicillin, 30 μg/ml kanamycin, 0.1 mM IPTG and 50 μg/ml X-Gal. Images were taken after incubation for 24 h and 48 h at 30°C.

## Results

### *L. monocytogenes* 10403S encodes six FtsW/RodA homologs

So far, FtsW and RodA proteins of the human pathogen *L. monocytogenes* have not been studied. FtsW and RodA are members of the SEDS (shape, elongation, division, sporulation) family of proteins and are multispanning membrane proteins with 8-10 transmembrane helices and a large extracellular loop (Fig. 1A). Using BLAST, six proteins with homology to the *B. subtilis* FtsW and RodA proteins could be identified in the genome of *L. monocytogenes* 10403S (Table 1). The protein encoded by *lmo0421* has the weakest homology to *B. subtilis* FtsW and RodA (Fig. S1, Table 1). *lmo0421* is part of the *sigC* operon, which is comprised of *lmo0422* encoding the PadR-like repressor LstR and *lmo0423* coding for the ECF-type sigma factor SigC (Fig. 1B). The *sigC* operon acts as a lineage II specific heat shock system (Zhang et al., 2005) and is therefore not encoded in all *L. monocytogenes* genomes. Due to the weak homology to FtsW and RodA and its absence in *L. monocytogenes* strains of lineage I and III, Lmo0421 was excluded from further analysis.

**Figure 1:**
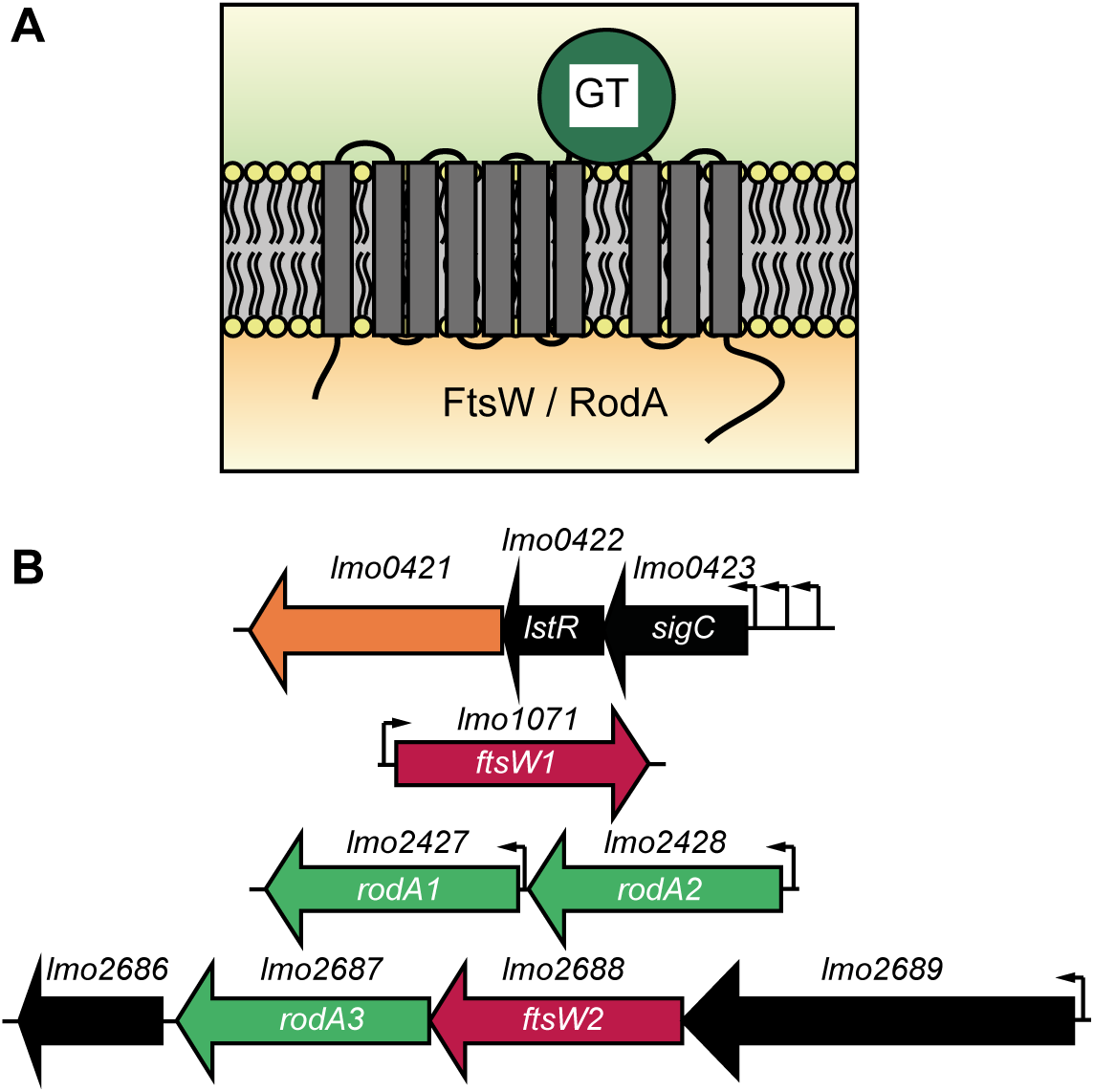
Topology of FtsW and RodA enzymes and their genomic arrangement in *L. monocytogenes*. (A) Topology models for FtsW and RodA enzymes of *L. monocytogenes* as predicted using the TMHMM v. 2.0 server (Sonnhammer et al., 1998). (B) Genomic organization of *ftsW/rodA* genes in *L. monocytogenes*. Arrowheads indicate the gene orientation. Small black arrows indicate promoters based on the operon structures described in Toledo-Arana *et al*. (Toledo-Arana et al., 2009). Three different promoters have been identified for the *sigC* operon (Zhang et al., 2005).

**Table 1:** Sequence homology between *B. subtilis* and *L. monocytogenes* FtsW/RodA proteins as determined by BLAST.

| ***L. monocytogenes* proteins** | ***B. subtilis* FtsW (403 aa)<sup>\*</sup>** | ***B. subtilis* RodA (393 aa)<sup>\*</sup>** |
| --- | --- | --- |
| **Lmo0421** | 46/128 aa (36%) | 77/350 aa (22%) |
| **(416 aa)** | 7e-14 | 4e-09 |
| **FtsW1 (Lmo1071)** | 195/403 aa (48%) | 100/333 aa (30%) |
| **(402 aa)** | 1e-122 | 7e-35 |
| **RodA1 (Lmo2427)** | 124/403 aa (31%) | 132/360 aa (37%) |
| **(391 aa)** | 9e-29 | 1e-57 |
| **RodA2 (Lmo2428)** | 112/374 aa (30%) | 159/402 aa (40%) |
| **(389 aa)** | 1e-33 | 1e-66 |
| **RodA3 (Lmo2687)** | 95/324 aa (29%) | 110/356 aa (31%) |
| **(369 aa)** | 3e-25 | 3e-44 |
| **FtsW2 (Lmo2688)** | 147/342 aa (43%) | 95/300 aa (32%) |
| **(376 aa)** | 5e-67 | 6e-22 |

<sup>\*</sup> *L. monocytogenes* proteins and their amino acid sizes (aa) are shown in bold in the left column and *B. subtilis* FtsW and RodA proteins used for the homology search are denoted in the top row. The number of amino acids, which were found to be identical between the respective *L. monocytogenes* and *B. subtilis* FtsW/RodA protein are denoted (left number) along with the amino acids that was utilized by the BLAST algorithm for this comparison (right number). The % identity is given in brackets and the corresponding e-values are indicated below.

The *L. monocytogenes* protein Lmo1071 is the closest homolog to *B. subtilis* FtsW with a sequence identity of 48% (Table 1). Furthermore, *L. monocytogenes lmo1071* and *B. subtilis ftsW* are found in the same chromosomal context. More specifically, *lmo1071* is located between genes *lmo1070*, which encodes a protein with homology to the *B. subtilis* YlaN protein, and *pycA* coding for the pyruvate carboxylase, which are also adjacent to *ftsW* in *B. subtilis*. This analysis suggests that gene *lmo1071* encodes the cell division protein FtsW. However, *L. monocytogenes* encodes a second protein, Lmo2688, that shares a higher degree of homology to the *B. subtilis* FtsW as compared to the *B. subtilis* RodA protein (Table 1). Due to these similarities and additional data presented in this study, we refer to Lmo1071 and Lmo2688 as FtsW1 and FtsW2, respectively.

The BLAST search with the *B. subtilis* RodA sequence as a query sequence yielded the *L. monocytogenes* protein Lmo2428 as the closest homolog with a sequence identity of 40% (Table 1). In addition to Lmo2428, two additional RodA homologs are present in *L. monocytogenes*, namely Lmo2427 and Lmo2687. As presented below, Lmo2427, Lmo2428 and Lmo2687 are likely bona-fide RodA homologs and were therefore renamed RodA1, RodA2 and RodA3, respectively. *rodA1* is located adjacent to *rodA2*, but despite their proximity, *rodA1* and *rodA2* are likely not transcribed as part of the same operon (Toledo-Arana et al., 2009). In contrast, *rodA3* and *ftsW2* are part of the four-gene operon *lmo2689-lmo2686*. Lmo2689 is similar to a Mg^2+^-type ATPase, whereas *lmo2686* encodes a protein of unknown function. Analysis of around 2000 genomes of *L. monocytogenes* strains presently available at NCBI revealed that the five FtsW/RodA homologs named here FtsW1, FtsW2, RodA1, RodA2 and RodA3 are conserved in the different strains with one exception; *L. monocytogenes* 4b strains of the sequence type ST2 lack homologs of RodA3 and FtsW2.

### *lmo1071* encodes FtsW1 and is essential for the survival of *L. monocytogenes*

The cell division protein FtsW is essential for growth in the Gram-negative and Gram-positive model organisms *E. coli* and *B. subtilis* (Boyle et al., 1997, Ikeda et al., 1989, Khattar et al., 1994, Kobayashi et al., 2003). Depletion of FtsW in these organisms leads to a block in cell division and formation of elongated cells (Boyle et al., 1997, Gamba et al., 2016). All our attempts to delete the *ftsW1* gene in *L. monocytogenes* 10403S remained unsuccessful, suggesting that FtsW1 is also essential for growth in *Listeria*. Next, strain 10403S Δ*ftsW1* i*ftsW1* was constructed, in which the expression of *ftsW1* is controlled by an IPTG-inducible promoter. While no difference in the growth was observed between the wildtype and FtsW1-depletion strain (likely due to leakiness of the inducible promoter) (Fig. 2A), cells depleted for FtsW1 were significantly elongated (Fig. 2B-C). Bacteria depleted for FtsW1 had a median cell length of 3.41±0.16 μm, while wildtype and 10403SΔ*ftsW1* i*ftsW1* bacteria grown in the presence of 1 mM IPTG had a median cell length of 1.85±0.16 μm and 1.93±0.07 μm, respectively (Fig. 2C). These data indicate that *lmo1071* encodes the cell division specific SEDS protein FtsW.

**Figure 2:**
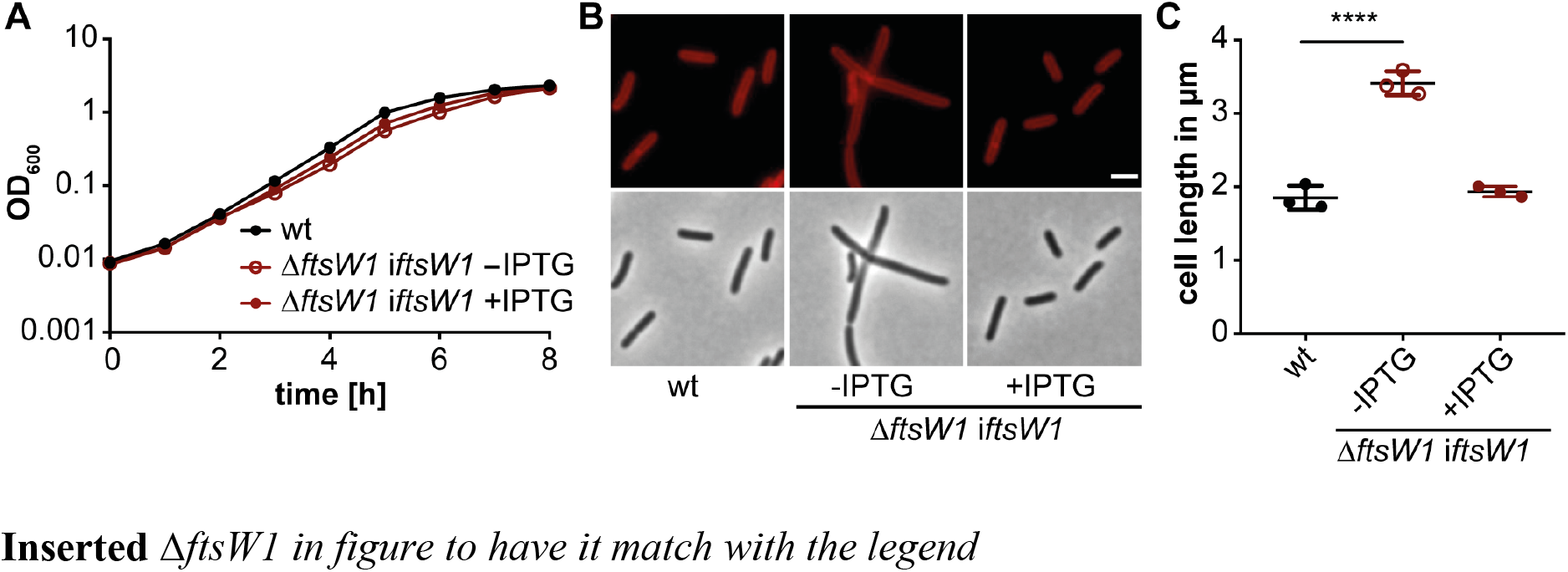
Depletion of FtsW1 results in cell elongation. (A) Growth of the *L. monocytogenes* FtsW1 depletion strain. Strain 10403SΔ*ftsW1* i*ftsW1* was grown overnight in BHI medium in the presence of 1 mM IPTG. The following day, the cells were washed and used to inoculate fresh BHI medium without IPTG and incubated at 37°C for 8-10h, diluted again and grown until the next morning. The depletion culture was used to inoculate BHI medium without IPTG or with 1 mM IPTG and the growth was monitored by determining OD_600_ readings at hourly intervals. Wild type *L. monocytogenes* strain 10403S (wt) was used as control. Averages and standard deviations from three independent experiments were plotted. Of note, due to the small standard deviations the actual error bars are not visible in the graph. (B) Microscopy analysis of 10403S (wt) and the *ftsW1* depletion strain grown in the presence or absence of IPTG. For depletion of FtsW1, strain 10403SΔ*ftsW1* i*ftsW1* was grown as described in panel A. Cultures of 10403S and 10403SΔ*ftsW1* i*ftsW1* were diluted 1:100 in fresh BHI medium (with 1 mM IPTG where indicated), grown for 3 h at 37°C and the cells subsequently stained with the membrane dye nile red and analyzed by fluorescence and phase contrast microscopy. The scale bar is 2 μm. (C) Cell lengths measurement of 10403S (wt) and the *ftsW1* depletion strain. The cell length of 300 cells per strain was measured and the median cell length calculated and plotted. Three independent experiments were performed, and the average and standard deviation of the median cell length plotted. For statistical analysis, a one-way ANOVA coupled with a Dunnett’s multiple comparisons test was performed (**** p≤0.0001).

### *L. monocytogenes* encodes a second FtsW protein

To our knowledge, all bacteria analyzed to date possess only one FtsW protein that is essential for cell survival. We identified a second potential FtsW protein, Lmo2688, in *L. monocytogenes*. In contrast to *ftsW1*, a *L. monocytogenes ftsW2* deletion strain could be constructed and no significant growth or cell morphology phenotypes could be observed for the Δ*ftsW2* deletion strain (Fig. S2). In a previous study, it was reported that the operon comprised of genes *lmo2689-lmo2686* is only minimally expressed when *L. monocytogenes* 10403S is grown in BHI broth (Lobel and Herskovits, 2016). We reasoned that if *ftsW2* does indeed code for a second FtsW protein, it should be possible to delete *ftsW1* in a strain in which *ftsW2* is artificially expressed from an IPTG-inducible promoter. Indeed, strain 10403SΔ*ftsW1* i*ftsW2* could be generated in the presence of IPTG. In contrast, we were unable to generate strain 10403SΔ*ftsW1* when any of the other FtsW/RodA homologs Lmo2427 (RodA1), Lmo2428 (RodA2) or Lmo2687 (RodA3) were expressed from the same IPTG-inducible promoter system. While prolonged depletion of FtsW2 in strain 10403SΔ*ftsW1* i*ftsW2* had again no impact on the growth, the cells were significantly elongated in the absence of the inducer compared to wildtype or bacteria grown in the presence of inducer (Fig. 3). These data strongly suggest that *ftsW2* encodes a second FtsW enzyme while the remaining three proteins Lmo2427, Lmo2428 and Lmo2687 likely function as RodA proteins.

**Fig. 3:**
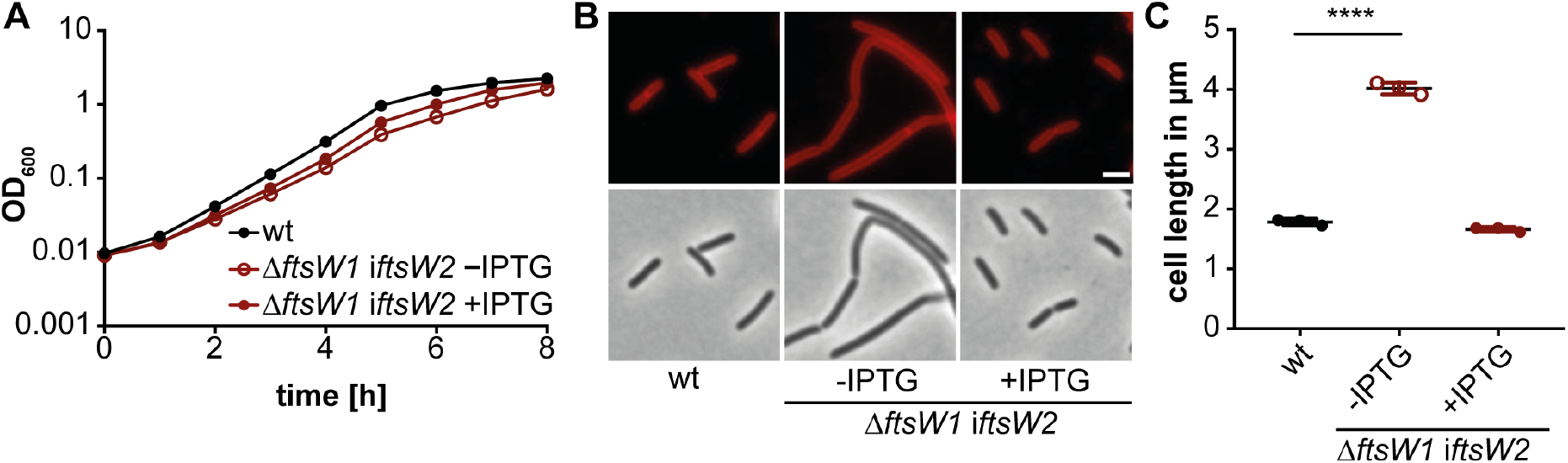
Overexpression of FtsW2 can compensate for the loss of FtsW1. (A) Growth comparison of strains 10403S (wt) and 10403SΔ*ftsW1* i*ftsW2*. Growth curves were performed as described in Figure 2. (B) Microscopy images of strains 10403S (wt) and 10403SΔ*ftsW1* i*ftsW2*. For depletion of FtsW2, strain 10403SΔ*ftsW1* i*ftsW2* was grown as described in Figure 2. The next morning, cells were diluted 1:100 and grown in BHI broth (with or without IPTG as indicated) until mid-logarithmic growth phase, stained with the membrane dye nile red and subjected to fluorescence and phase contrast microscopy analysis. Scale bar is 2 μm. (C) Cell lengths measurements of 10403S (wt) and 10403SΔ*ftsW1* i*ftsW2*. The cell length of 300 cells per strain was measured and the median cell length calculated and plotted. Three independent experiments were performed, and the average and standard deviation of the median cell length plotted. For statistical analysis, a one-way ANOVA coupled with a Dunnett’s multiple comparisons test was performed (**** p≤0.0001).

### *L. monocytogenes* encodes three RodA homologs

We were able to assign roles for two of the FtsW/RodA homologs as FtsW-like proteins. However, *L. monocytogenes* encodes three additional homologs, which show a higher similarity to the *B. subtilis* RodA protein as compared to the *B. subtilis* FtsW protein (Table 1). As described above, expression of none of the enzymes Lmo2427, Lmo2428 or Lmo2687 was able to rescue the growth of an *ftsW1* deletion strain, indicating that these enzymes likely function as RodA proteins in *L. monocytogenes* and hence they were renamed RodA1, RodA2 and RodA3, respectively. All attempts to construct a *rodA1-3* triple mutant failed further corroborating that these proteins function as RodA proteins and at least one of them needs to be present for cell viability. To determine whether the different RodA homologs have distinct functions or whether they are merely duplications, single and double mutant strains were generated. No significant differences with regards to growth and cell length could be observed between the wildtype strain 10403S and single *rodA1, rodA2* or *rodA3* deletion strains (Fig. S3). Similar observations were made with the *rodA* double mutant strains 10403S Δ*rodA1*Δ*rodA2* and 10403SΔ*rodA2*Δ*rodA3* (Fig. 4). However, cells lacking RodA1 and RodA3 were shorter (1.3±0.03 μm) compared to wildtype cells (1.9±0.06 μm, Fig. 4B-C), indicating that either RodA1 or RodA3 needs to be present for *L. monocytogenes* to maintain its rod shape. RodA3 is part of the *lmo2689-lmo2686* operon that is only minimally expressed when *L. monocytogenes* is grown in BHI broth (Lobel and Herskovits, 2016). The fact that we observe differences in cell morphology between a *rodA1* single and the *rodA1/rodA3* double mutant, suggests that the function of the two proteins could be additive or that *rodA3* expression, which is only minimally expressed in a wildtype strain under standard laboratory growth conditions, might increase upon deletion of *rodA1*. To test if *rodA3* expression is increased in the absence of other RodA proteins, we fused the promoter upstream of *lmo2689* and driving *rodA3* expression to *lacZ* and inserted this fusion into the chromosome of wildtype 10403S, the *rodA1* and the *rodA1/rodA2* deletion strains. The promoter activity was indeed 1.5 – to 2-fold higher in the *rodA1* and *rodA1/rodA2* mutant strains as compared to the wildtype, as assessed by the increase in the β-galactosidase activity (Fig. 4G). This result indicates that expression of the *lmo2689-lmo2686* operon, which encodes FtsW2 and RodA3, is induced in the absence of RodA1, suggesting a coordination of the expression of the different RodA homologs.

**Fig. 4:**
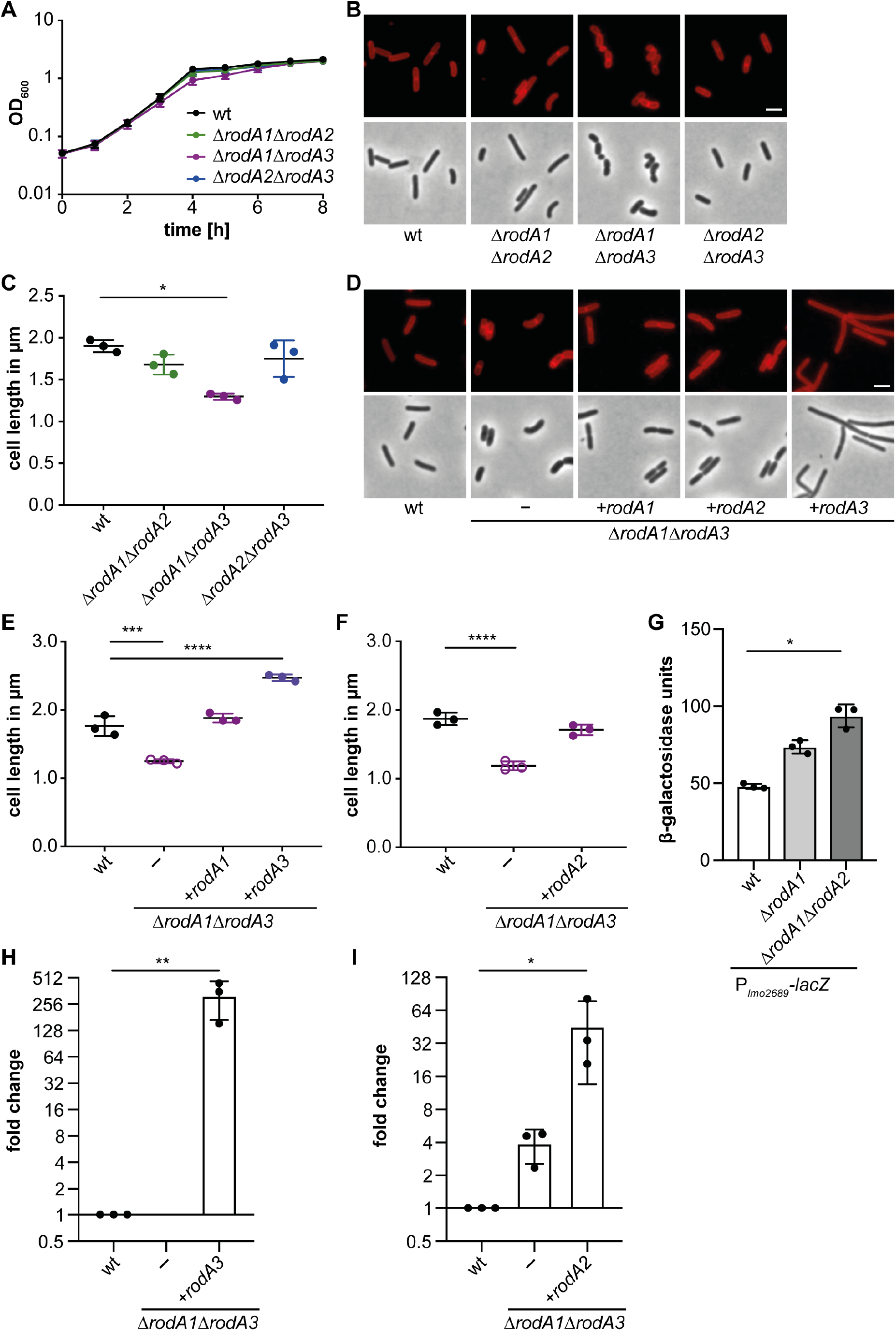
RodA1 and RodA3 are essential to maintain the rod shape of *L. monocytogenes*. (A) Growth of *L. monocytogenes* 10403S (wt), 10403SΔ*rodA1*Δ*rodA2*, 10403SΔ*rodA1*Δ*rodA3* and 10403SΔ*rodA2*Δ*rodA3* strains in BHI broth at 37°C. (B) Microscopy images of wt and mutant *L. monocytogenes* strains. Cells from the strains described in panel A were stained with the membrane dye nile red and analyzed by fluorescence and phase contrast microscopy. Scale bar is 2 μm. (C) Cell lengths measurements of wt and mutant *L. monocytogenes* strains. The cell length of 300 cells per strain was measured and the median cell length calculated and plotted. Three independent experiments were performed, and the average and standard deviation of the median cell length plotted. For statistical analysis, one-way ANOVA t-tests for multiple comparisons were used (* p≤0.05). (D) Microscopy images of wildtype and mutant *L. monocytogenes* strains. Cells from 10403S (wt), 10403S.Δ*rodA1*Δ*rodA3* (-) and 10403SΔ*rodA1*Δ*rodA3* expressing *rodA1, rodA2* or *rodA3* from an IPTG-inducible promoter were stained with the membrane dye nile red and analyzed by fluorescence and phase contrast microscopy. Scale bar is 2 Δm. (E-F) Cell length analysis of 10403SΔ*rodA1* Δ*rodA3* complementation strains. Analysis was performed as described in panel C. (G) β-galactosidase assay using *L. monocytogenes* strains 10403S (wt), 10403SΔ*rodA1* and 10403SΔ*rodA1-2* carrying a *P_lmo2689-_lacZ* fusion construct. Averages and standard deviations of three independent experiments were plotted. For statistical analysis, a one-way ANOVA coupled with a Dunnett’s multiple comparisons test was used (* p≤0.05). (H) Analysis of *rodA3* expression by qRT-PCR. RNA was isolated from strains 10403S (wt), 10403SΔ*rodA1*Δ*rodA3* (-) and 10403SΔ*rodA1*Δ*rodA3* i*rodA3* grown in the presence of IPTG. Expression of *rodA3* was normalized to the expression of *gyrB* and fold changes calculated using the ΔΔCt method. Averages and standard deviations of three independent experiments were plotted. For statistical analysis, a one-way ANOVA coupled with a Dunnett’s multiple comparisons test was performed (** p≤0.01). (I) Analysis of *rodA2* expression using qRT-PCR. Same as panel (H) but using strains 10403S, 10403SΔ*rodA1* Δ*rodA3* (-) and 10403SΔ*rodA1* Δ*rodA3* i*rodA2* grown in the presence of IPTG. Averages and standard deviations of three independent experiments were plotted. For statistical analysis, a one-way ANOVA coupled with a Dunnett’s multiple comparisons test was performed (* p≤0.05).

To confirm that the decrease in cell length of the double mutant strain 10403S Δ*rodA1*Δ*rodA3* depends on the absence of RodA1 and RodA3, complementation strains with IPTG-inducible expression of *rodA1* or *rodA3* were constructed. Expression of RodA1 restored the cell length to 1.84±0.1 μm, which is comparable to the cell length of wildtype cells (1.76±0.15 μm, Fig. 4E). On the other hand, expression of *rodA3* from an ectopic locus and IPTG-inducible promoter in strain 10403SΔ*rodA1*Δ*rodA3* led to the formation of longer cells with an average cell length of 2.47±0.01 μm (Fig. 4D-E). These results indicate that induction of RodA3 from the IPTG-inducible promoter likely results in an overproduction of the protein as compared to the expression from the native promoter. To confirm that *rodA3* expression is increased when expressed from the IPTG-inducible promoter as compared to its native promoter, *rodA3* transcript levels were assessed by qRT-PCR in the wildtype strain, the 10403SΔ*rodA1*Δ*rodA3* deletion strain and strain 10403SΔ*rodA1*Δ*rodA3* i*rodA3* grown in the presence of IPTG (Fig. 4H). Significant higher *rodA3* transcript levels were detected in the inducible strain in the presence of IPTG as compared to the wildtype strain. Similarly, expression of *rodA3* from an IPTG-inducible promoter in wildtype 10403S led to the formation of elongated cells with an average cell length of 2.49±0.08 μm, whereas additional expression of *rodA1* or *rodA2* using the same inducible system had no impact on the cell length of 10403S (Fig. S4). These results highlight that in particular fine-tuning of RodA3 production is essential for cell-length determination in *L. monocytogenes*.

The observation that the *rodA1/rodA3* double mutant forms shorter cells suggests that RodA2 is not sufficient to maintain the cell length of *L. monocytogenes*. There are several possible explanations for this: RodA2 might have a reduced activity as compared to RodA1 or RodA3. RodA2 might have a function that is different from RodA1 and RodA3 or the protein levels of RodA2 might be sufficient to maintain cell viability but too low to maintain the rod shape. To investigate this further, a strain was constructed which lacks *rodA1* and *rodA3*, but carries pIMK3-*rodA2* to allow for IPTG-inducible expression of *rodA2* in addition to the expression of *rodA2* from its native locus (10403SΔ*rodA1*Δ*rodA3* i*rodA2*). In the absence of the inducer, the cells had a median cell length of 1.2±0.03 μm (data not shown). However, the cell length of strain 10403SΔ*rodA1*Δ*rodA3* i*rodA2* increased to 1.71±0.8 μm when the strain was grown in the presence of IPTG (Fig. 4F). Therefore, additional expression of *rodA2*, which was verified by qRT-PCR (Fig. 4I), can partially complement the cell length phenotype of the *rodA1/rodA3* deletion strain, suggesting that RodA2 has a similar function as RodA1 and RodA3, but that it has either a lower activity or is not expressed in sufficient amounts from its native promoter for proper cell length maintenance.

As stated above, several attempts to construct a strain inactivated for all three RodA homologs remained unsuccessful, suggesting that at least one of the proteins RodA1, RodA2 or RodA3 needs to be present for the viability of *L. monocytogenes*. The results presented so far indicate that RodA1 is the most important RodA homolog considering that RodA2 alone is not sufficient to maintain the rod shape and that RodA3 is only minimally expressed under standard laboratory conditions (Lobel and Herskovits, 2016) and also lacking in *L. monocytogenes* 4b strains of the sequence type ST2. To understand the impact of RodA enzymes on cell growth and cell division in *L. monocytogenes*, a strain was constructed which lacks all three *rodA* genes from its genome, but harbors pIMK3-*rodA1* to enable IPTG-inducible expression of RodA1. Prolonged depletion of RodA1 in strain 10403SΔ*rodA1-3* i*rodA1* led to a growth defect that was not seen when the strain was grown in the presence of the inducer (Fig. 5A). However, the depletion was not efficient enough to see a complete growth inhibition, which would be expected for a strain lacking all three RodA homologs. This is likely caused by residual *rodA1* expression from the inducible promoter even in the absence of IPTG. Consistent with this notion, even after prolonged depletion, *rodA1* transcripts could still be detected in strain 10403SΔ*rodA1-3* i*rodA1* as assessed by qRT-PCR (Fig. S5). However, cells of the *L. monocytogenes* strain 10403SΔ*rodA1-3* i*rodA1* that were grown without IPTG were significantly shorter with a cell length of 1.18±0.08 μm as compared to cells of the double mutant 10403SΔ*rodA1*Δ*rodA3* or the wildtype strain 10403S (Fig. 5B, D). Interestingly, different cell morphologies could be observed for strain 10403SΔ*rodA1-3* i*rodA1* after prolonged RodA1 depletion (Fig. 5C). The placement of the division septum was affected in some cells and daughter cells of different size or cells with two septa were observed (Fig. 5C). These morphological defects could be complemented, and the cell length increased to 1.95±0.04 μm upon addition of IPTG and expression of RodA1 (Fig. 5D). These data highlight that RodA1 alone is sufficient to maintain the cell shape of *L. monocytogenes*.

**Figure 5:**
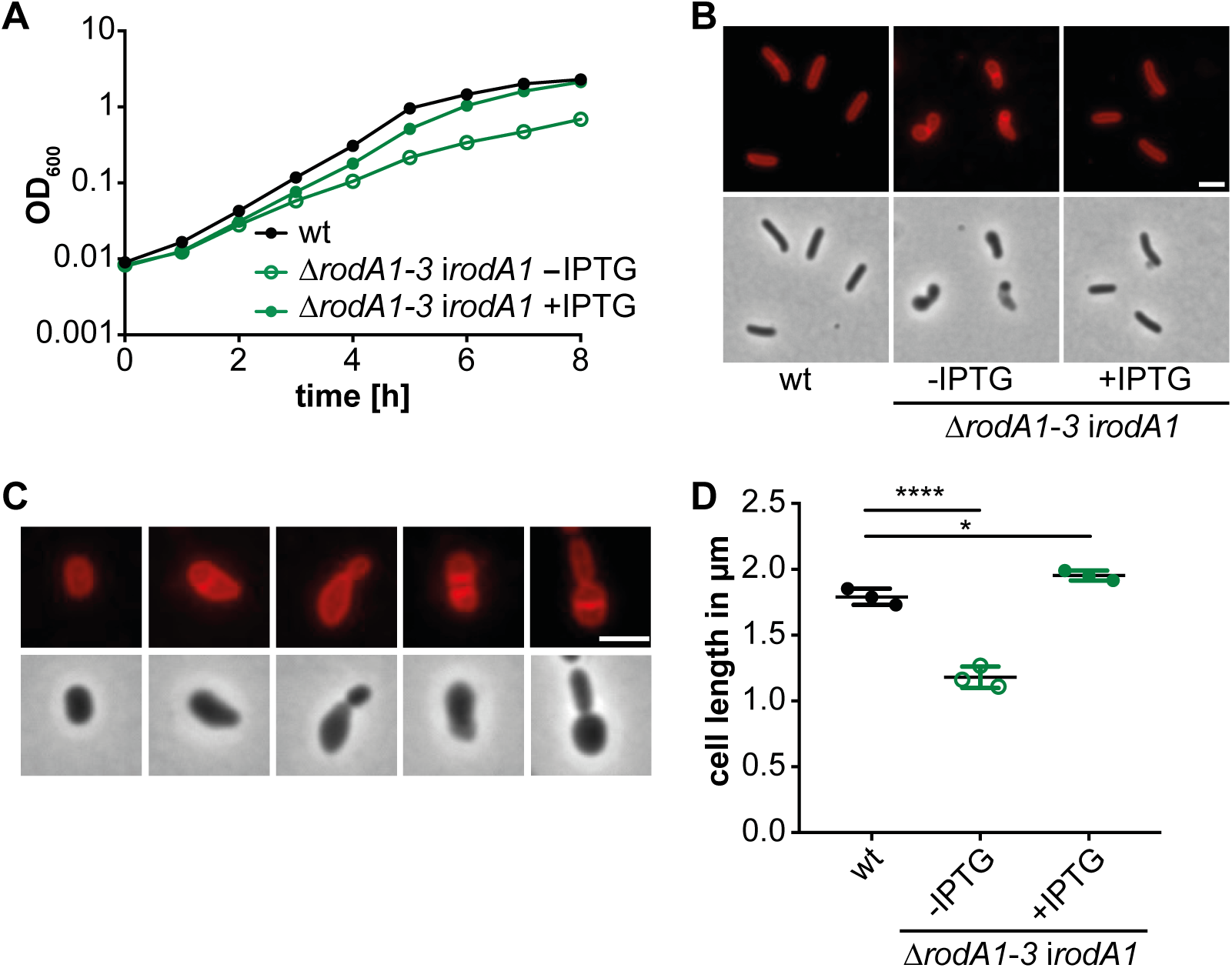
Absence of all three RodA proteins leads to a drastic cell shortening. (A) Growth of *L. monocytogenes* in the absence of all three RodA proteins. Growth curves were performed as described in Figure 2 using strain 10403S and strain 10403SΔ*rodA1-3* i*rodA1* grown in the presence of absence of IPTG. (B and C) Microscopy analysis of strain 10403SΔ*rodA1-3* i*rodA1*. For depletion of RodA1, strain 10403SΔ*rodA1-3* i*rodA1* was grown in the same way as described for the FtsW1 depletion in Figure 2. (B) Wildtype 10403S and depleted 10403SΔ*rodA1-3* i*rodA1* cultures were diluted 1:100, grown for 3 h at 37°C (where indicated in the presence of 1 mM IPTG), stained with nile red and phase contrast and fluorescence microscopy images were taken. Scale bar is 2 μm. (C) Microscopy images showing examples of different cell morphologies observed for strain 10403S Δ*rodA1-irodA1* depleted for RodA1. Scale bar is 2 μm. (D) Cell lengths measurements of 10403S (wt) and strain 10403SΔ*rodA1-3* i*rodA1*. The cell length of 300 cells per strain was measured. Three independent experiments were performed, and the average and standard deviation of the median cell length plotted. For statistical analysis, a one-way ANOVA coupled with a Dunnett’s multiple comparisons test was performed (* p≤0.05, **** p<0.0001).

### Decreased moenomycin and lysozyme resistance in the absence of RodA homologs

Next, we wondered whether the absence of FtsW or RodA proteins affects the resistance of *L. monocytogenes* towards the antibiotics penicillin, bacitracin and moenomycin, which target different steps in the peptidoglycan biosynthesis process. Penicillin binds to the transpeptidase domain of PBPs and inhibits their function, leading to a reduced crosslinking of the peptidoglycan (Nakagawa et al., 1979, Korsak et al., 2010). Bacitracin inhibits the dephosphorylation of the bactoprenol carrier leading to a block in lipid II synthesis (Stone and Strominger, 1971). The phosphoglycolipid antibiotic moenomycin inhibits the glycosyltransferase activity of bifunctional PBPs and thereby prevents the polymerization of the glycan chain (van Heijenoort et al., 1987).

No significant differences could be observed in terms of resistance against penicillin, bacitracin or moenomycin for the FtsW1 depletion strain 10403SΔ*ftsW1* i*ftsW1*. This is presumably due to basal level expression of *ftsW1* even in the absence of the inducer. Simultaneous deletion of *rodA1* and *rodA3* resulted in a slight decrease in the MIC for penicillin, however, this difference was not significant (Fig. 6A). On the other hand, strain 10403SΔ*rodA1*Δ*rodA3* was 2-4-fold more sensitive to the antibiotic bacitracin (Fig. 6B). This phenotype could be complemented by expressing either RodA1, RodA2 or RodA3 from an IPTG-inducible promoter (Fig. 6B).

**Figure 6:**
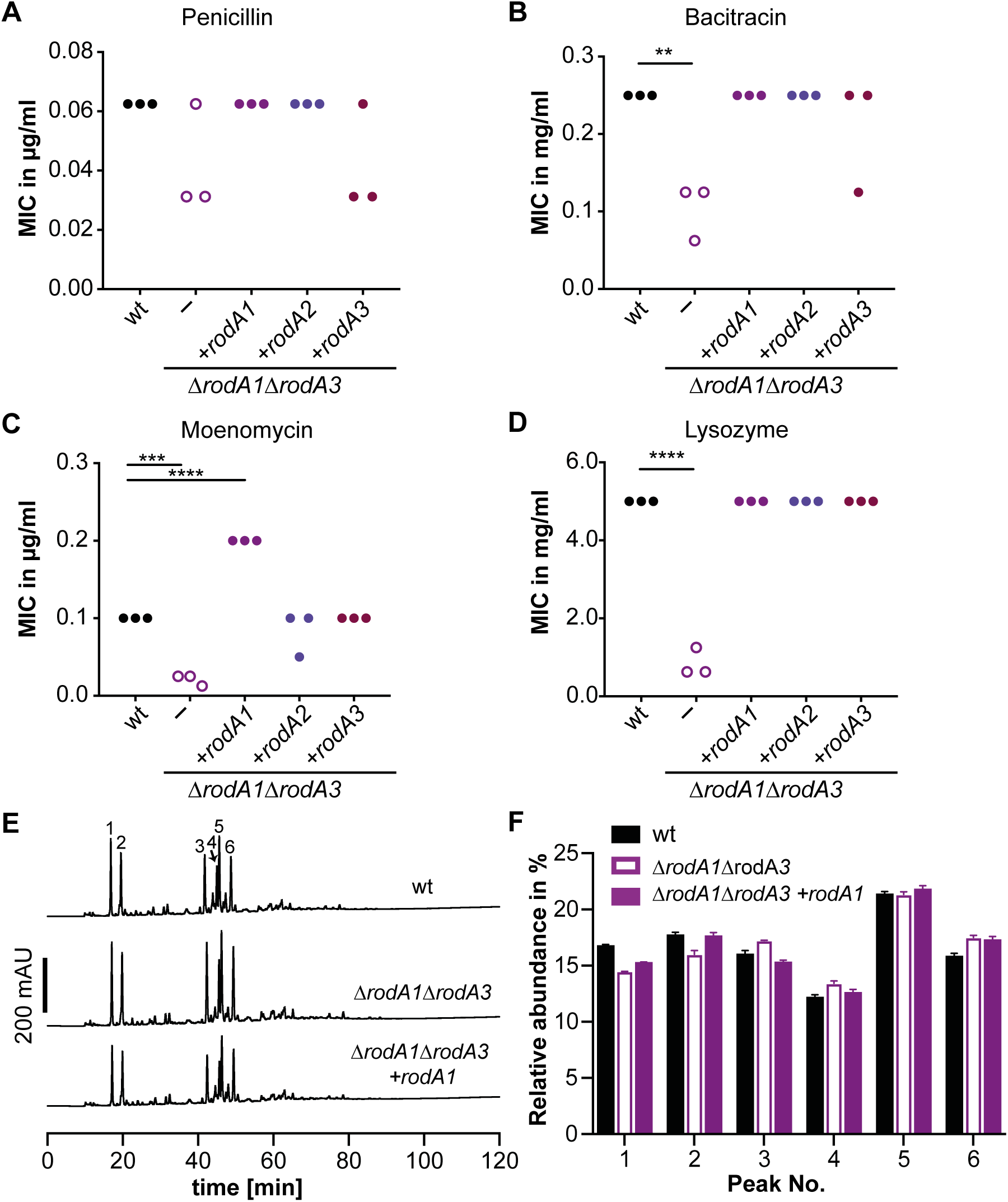
Impact of *rodA1* and *rodA3* deletion on the antibiotic and lysozyme resistance. The minimal inhibitory concentrations for the antibiotics penicillin G (A), bacitracin A (B), moenomycin (C) and lysozyme (D) were determined for the wildtype *L. monocytogenes* strain 10403S, the Δ*rodA1*Δ*rodA3* deletion strain and the complementation strains 10403SΔ*rodA1*Δ*rodA3* i*rodA1*, 10403SΔ*rodA1*Δ*rodA3* i*rodA2* and 10403SΔ*rodA1*Δ*rodA3* i*rodA3* using a microbroth dilution assay. The complementation strains were grown in the presence of 1 mM IPTG. The result of three biological replicates are shown in A-D. For statistical analysis, a one-way ANOVA coupled with a Dunnett’s multiple comparisons test was used (** p≤0.01, *** p≤0.001, **** p≤0.0001). (E) HPLC analysis of the muropeptide composition of 10403S (wt), 10403SΔ*rodA1*Δ*rodA3* and 10403SΔ*rodA1*Δ*rodA3* i*rodA1* (grown in the presence of IPTG to induce expression of RodA1). The major muropeptide peaks are numbered 1-6 as previously described (Rismondo et al., 2015, Burke et al., 2014). (F) Relative abundance of muropeptide peaks 1-6 in peptidoglycan isolated from strains 10403S (wt), 10403SΔ*rodA1*Δ*rodA3* and 10403SΔ*rodA1* Δ*rodA3* i*rodA1*. Average values and standard deviations were calculated from three independent peptidoglycan extractions.

As described above, moenomycin inhibits the transglycosylase activity of PBPs leading to a decreased activity of these enzymes. In the absence of RodA1 and RodA3, cells are more susceptible to a reduced activity of PBPs manifesting in a 4-fold reduced resistance to moenomycin (Fig 6C). Induction of RodA1 expression in strain 10403SΔ*rodA1*Δ*rodA3* resulted in a significantly higher resistance to moenomycin as compared to the wildtype strain 10403S and expression of RodA2 or RodA3, led to partial or complete complementation of the moenomycin sensitivity (Fig. 6C).

Moreover, resistance to lysozyme, an enzyme that cleaves the linkage between N-acetyl muramic acid and N-acetylglucosamine residues of the peptidoglycan, was drastically decreased in strain 10403SΔ*rodA1*Δ*rodA3* and could be fully restored by expression of RodA1, RodA2 or RodA3 (Fig. 6D). Lysozyme resistance in *L. monocytogenes* is mainly accomplished by two modifications of the peptidoglycan; deacetylation of N-acetylglucosamine residues by PgdA or O-acetylation of N-acetylmuramic acid residues by OatA (Boneca et al., 2007, Aubry et al., 2011). To determine whether the activity of PgdA is changed in the absence of RodA1 and RodA3, peptidoglycan was purified from the 10403SΔ*rodA1*Δ*rodA3* mutant strain, digested with mutanolysin and the resulting muropeptides analyzed by HPLC. Peptidoglycan samples isolated from the wildtype strain 10403S and the complementation strain 10403SΔ*rodA1*Δ*rodA3* i*rodA1*, that had been grown in the presence of IPTG, were analyzed as controls (Fig. 6E). The main muropeptide peaks were assigned as described previously (Rismondo et al., 2015, Burke et al., 2014). Peaks 1 and 2 correspond to the acetylated and deacetylated monomeric muropeptides, respectively, whereas peak 3 and peaks 4-6 are acetylated and deacetylated muropeptide dimers, respectively. Deletion of *rodA1* and *rodA3* led to a reduction of both monomeric muropeptides and therefore to an increase in crosslinked peptidoglycan fragments by approximately 2% as compared to the wildtype strain 10403S, in which 65% of the peptidoglycan was cross-linked (Fig. 6F). However, no significant difference with regards to the deacetylated muropeptides could be observed between wildtype 10403S, strain 10403SΔ*rodA1*Δ*rodA3* and the 10403SΔ*rodA1*Δ*rodA3* i*rodA1* complementation strain (Fig. 6F). These results suggest that the lysozyme sensitivity phenotype of strain 10403SΔ*rodA1*Δ*rodA3* is not caused by changes in the peptidoglycan deacetylation, but rather due to general defects in the peptidoglycan structure.

### Cell wall-acting antibiotics induce the promoter of *lmo2689*

The operon *lmo2689-lmo2686*, that contains the genes encoding FtsW2 and RodA3, is only minimally expressed under standard laboratory conditions (Lobel and Herskovits, 2016). A genome-wide transcriptional analysis performed in *L. monocytogenes* strain LO28 has shown that *lmo2687, lmo2688* and *lmo2689* are part of the CesR regulon (Nielsen et al., 2012). The cephalosporin sensitivity response regulator CesR is part of the CesRK two-component system that regulates the transcription of several cell envelope-related genes in response to changes in cell wall integrity, such as caused by the presence of cell wall-acting antibiotics or alcohols including ethanol (Gottschalk et al., 2008, Kallipolitis et al., 2003, Nielsen et al., 2012). Therefore, we next used the *lmo2689 promoter-lacZ* fusion described above, to assess if expression of the *lmo2689-lmo2686* operon is induced in the presence of antibiotics that target different processes of the PG biosynthesis or ethanol. Indeed, increased β-galactosidase activity could be measured for cells that had been grown in the presence of sub-inhibitory concentrations of the β-lactam antibiotics ampicillin, penicillin and cefuroxime and the phosphoglycolipid moenomycin (Fig. 7A). In contrast, no increase in β-galactosidase activity could be detected upon addition of vancomycin, lysozyme or ethanol as compared to untreated control cells (Fig. 7A). We also tested whether the presence of MgSO4 or EDTA has an impact on the *lmo2689* promoter activity since *lmo2689* encodes a putative Mg^2+^-type ATPase. However, the β-galactosidase activity of cells grown in the presence of MgSO4 or EDTA was comparable to the β-galactosidase activity seen for untreated cells (Fig. 7A). These results indicate that the expression of *ftsW2* and *rodA3*, that are part of the *lmo2689-lmo2686* operon, are induced in the presence of various cell wall-acting antibiotics, suggesting that FtsW2 and RodA3 might be important for the intrinsic resistance of *L. monocytogenes* against these antibiotics. However, no significant differences in MICs for penicillin and moenomycin could be observed between wildtype 10403S, the *ftsW2* or *rodA3* single mutant strains or the *ftsW2/rodA3* double mutant (Fig. S6). However, there was a slight reduction in the resistance of the *rodA3* single mutant against cefuroxime as compared to the wildtype (Fig. S6). To further assess whether there is a difference in the cefuroxime resistance between the *L. monocytogenes* wildtype strain 10403S and the *rodA1, rodA2* and *rodA3* single mutant strains, dilutions of overnight cultures were spotted on BHI agar plates with or without 1 μg/ml cefuroxime. Deletion of *rodA1* or *rodA2* results in a slightly reduced ability of these strains to grow on BHI plates supplemented with 1 μg/ml cefuroxime as compared to the wildtype 10403S strain (Fig. 7B). However, deletion of *rodA3* leads to a stronger reduction of growth on BHI plates containing 1 μg/ml cefuroxime as compared to the *rodA1* and *rodA2* single mutants (Fig. 7B). Interestingly, overexpression of RodA3, but not of RodA1 or RodA2, also resulted in decreased resistance towards cefuroxime as compared to the wildtype strain 10403S (Fig. 7C). Our results therefore suggest that *L. monocytogenes* induces the expression of *rodA3* and *ftsW2* in the presence of β-lactam antibiotics and moenomycin to compensate for the inhibition of the glycosyltransferase and transpeptidase activity of PBPs. In particular RodA3 seems to play an important function for the intrinsic cephalosporine resistance in *L. monocytogenes* and its expression needs to be finely balanced as both its absence as well as increased expression have detrimental effects on resistance against this antibiotic.

**Figure 7:**
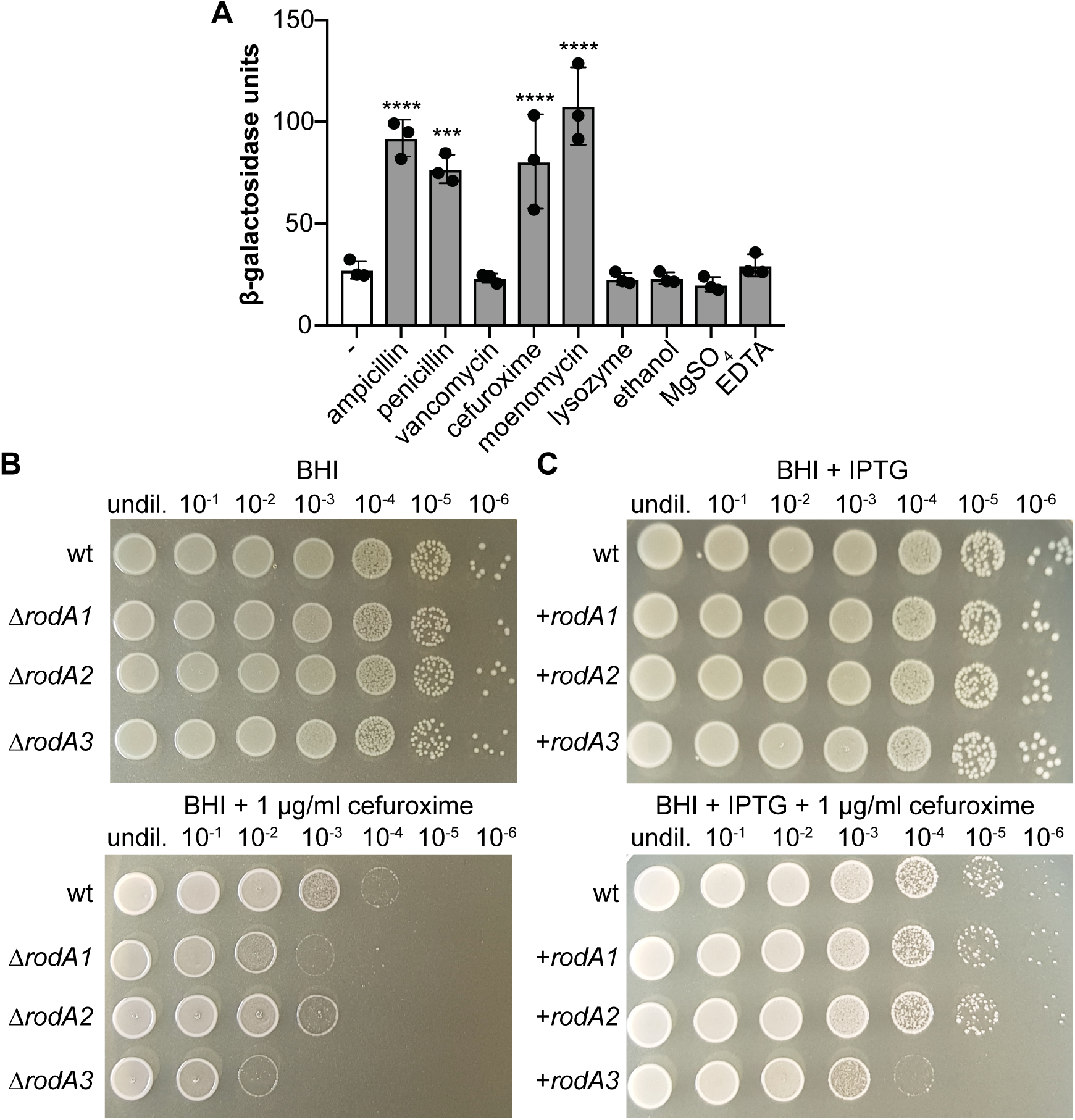
Cell wall-acting antibiotics increase the *lmo2689* promoter activity. (A) Bacteria from mid-logarithmic cultures of strain 10403S pPL3e-P*_lmo2689-_lacZ* were exposed for 2 h at 37°C to different stressors. The activity of the *lmo2689* promoter was subsequently determined by performing β-galactosidase activity assays as described in the method section. Bacteria that had been grown in the absence of a stressor were included as negative (-) control. The averages of the β-galactosidase activity units and standard deviations from three independent experiments were plotted. For statistical analysis, a one-way ANOVA coupled with a Dunnett’s multiple comparisons test was used (*** p≤0.001, **** p≤0.0001). (B) Absence and overexpression of RodA3 results in decreased cefuroxime resistance. Dilutions of overnight cultures of strains (B) 10403S (wt), 10403SΔ*rodA1*, 10403SΔ*rodA2* and 10403SΔ*rodA3* and (C) 10403S (wt), 10403S pIMK3-*rodA1*, 10403S pIMK3-*rodA2* and 10403S pIMK3-*rodA3* were spotted on BHI agar and BHI agar containing 1 μg/ml cefuroxime and incubated for 24 h at 37°C. BHI agar plates were supplemented with 1 mM IPTG for cefuroxime resistance assays shown in panel C. A representative result from three independent experiments is shown.

### FtsW and RodA proteins interact with class B PBPs

Previous studies have shown that members of the SEDS protein family act together with a cognate class B PBP to synthesize and crosslink peptidoglycan (Henriques et al., 1998, Wei et al., 2003, Reichmann et al., 2019, Gamba et al., 2009, Daniel et al., 1996). *L. monocytogenes* encodes three class B PBPs, namely PBP B1, PBP B2 and PBP B3. To identify potential protein-protein interactions between the *L. monocytogenes* class B PBPs and the FtsW/RodA proteins, a bacterial adenylate cyclase two-hybrid (BACTH) analysis was performed. Interactions were detected between the three *L. monocytogenes* PBP B1, PBP B2 and PBP B3 and all FtsW and RodA homologs (Fig. 8). These results suggest that the FtsW and RodA proteins also form a complex with class B PBPs in *L. monocytogenes*. However, using this bacterial two hybrid approach it was not possible to determine specific SEDS protein and class B PBP pairs.

**Figure 8:**
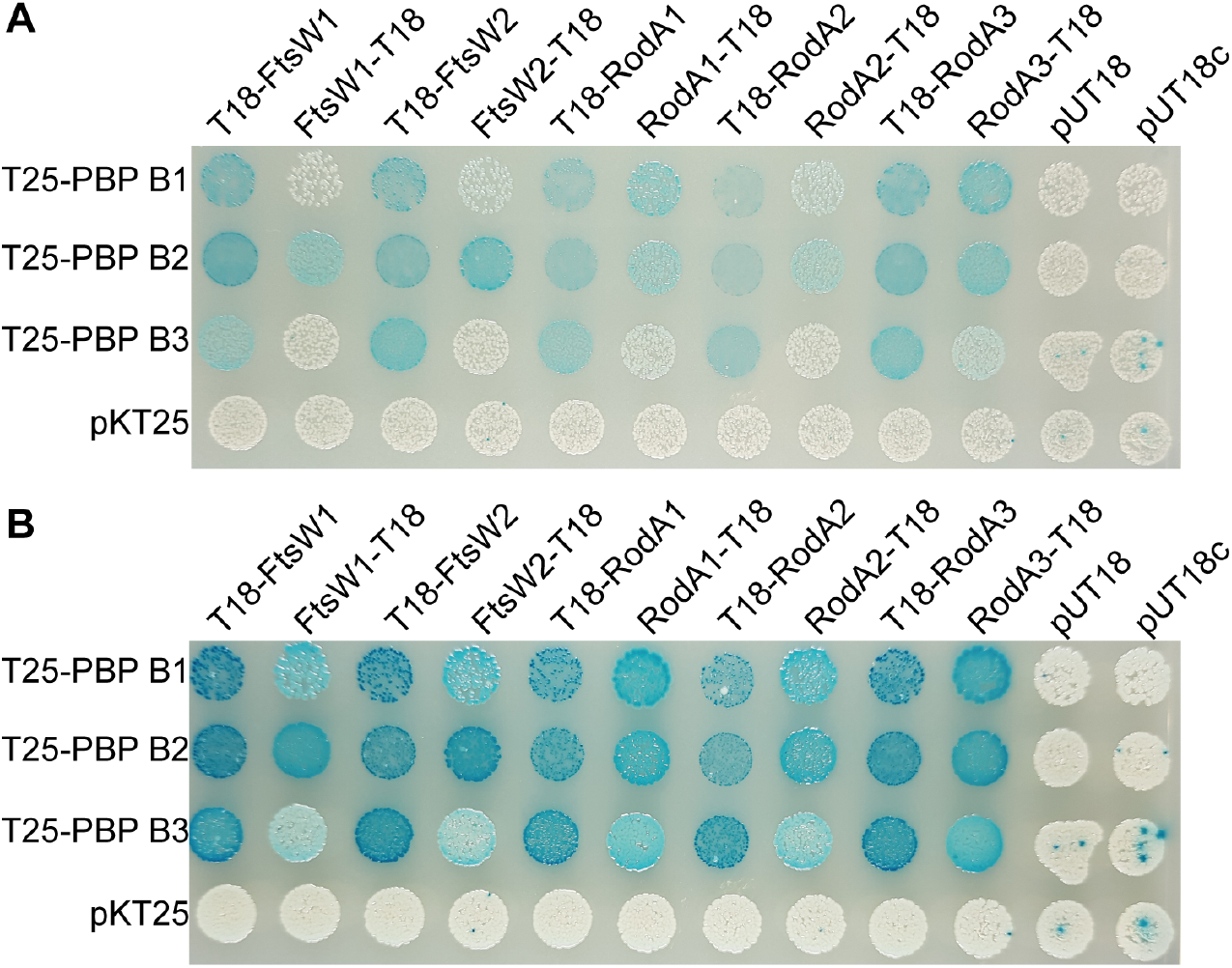
FtsW and RodA enzymes interact with class B PBPs. (A-B) FtsW/RodA and class B PBPs were fused to the T18 and T25-fragments of the *Bordetella pertussis* adenylate cyclase and co-transformed into the bacterial two-hybrid strain BTH101. Co-transformations with pKT25, pUT18 and pUT18c were used as negative controls. The plates were photographed after incubation at 30°C for (A) 24 h or (B) 48 h. Representative images from four independent experiments are shown.

## Discussion

Bacterial cell elongation and cell division need to be tightly regulated to maintain the cell shape. This is accomplished by two multiprotein complexes, the elongasome and the divisome, which are coordinated by the actin homolog MreB and the tubulin homolog FtsZ, respectively (Carballido-Lopez and Formstone, 2007, Typas et al., 2012, Jones et al., 2001, Errington and Wu, 2017, den Blaauwen, 2018). The SEDS protein FtsW is part of the divisome and essential for growth as shown for many bacteria including *E. coli, B. subtilis* and *S. aureus* (Boyle et al., 1997, Ikeda et al., 1989, Khattar et al., 1994, Kobayashi et al., 2003, Reichmann et al., 2019). Our experiments suggested that FtsW1 is also essential in *L. monocytogenes*, however, a second FtsW protein, FtsW2, can compensate for the loss of FtsW1 if it is expressed from an inducible promoter. FtsW2 is encoded in the *lmo2689-lmo2686* operon that appears to be only minimally expressed when *L. monocytogenes* 10403S is grown under standard laboratory conditions (Lobel and Herskovits, 2016). The expression of the *lmo2689-lmo2686* operon is regulated by the two-component system CesRK that is activated by cell envelope stress (Kallipolitis et al., 2003, Gottschalk et al., 2008, Nielsen et al., 2012). Using an *L. monocytogenes* strain carrying a *P_lmo2689-_lacZ* promoter fusion, we could detect increased β-galactosidase activity after incubation with sub-inhibitory concentrations of different β-lactam antibiotics including penicillin, cefuroxime, and moenomycin. However, the expression of the *lmo2689-lmo2686* operon was not induced by other cell wall-targeting antibiotics such as vancomycin or the hydrolase lysozyme. This suggests that inhibition of the glycosyltransferase or transpeptidase activity of PBPs leads to activation of the *lmo2689-lmo2686* operon, and hence, to the expression of *ftsW2* as well as *rodA3*.

The rod-shape determining protein RodA is part of the elongation machinery. The data presented in this study suggest that *L. monocytogenes* encodes not one but three RodA proteins and depletion of the three RodA enzymes leads to a decreased cell length (Fig. 5). Simultaneous deletion of *rodA1* and *rodA3* already results in the formation of shorter cells, whereas cells of strains deleted for *rodA1/rodA2* or *rodA2/rodA3* have a cell length that is comparable to the wildtype strain 10403S. Taking into consideration that *rodA3* is only minimally expressed under standard laboratory growth conditions in *L. monocytogenes* 10403S (Lobel and Herskovits, 2016), the results presented in this study suggest that *rodA3* expression gets induced upon inactivation of RodA1, since we observed morphological differences between the *rodA1* single and the *rodA1/3* double mutant strains. Indeed, β-galactosidase assays confirmed that deletion of *rodA1* or *rodA1/2* increases the activity of the promoter from which *rodA3* is expressed. The data presented in this study also indicate that RodA1 is the “main” RodA enzyme in *L. monocytogenes* as no significant phenotypic changes with regards to growth and cell division could be observed as long as RodA1 was present. On the other hand, RodA2 was only able to compensate for the loss of RodA1 and RodA3, when overproduced from an inducible promoter. Interestingly, cells of strains 10403S and 10403SΔ*rodA1*Δ*rodA3* in which *rodA3* is overexpressed from an ectopic locus have an increased cell length as compared to the wildtype. An explanation for this could be that elevated levels of RodA3 lead to the depletion of proteins needed at the cell division site, resulting in an extended synthesis of PG on the lateral wall. Another possibility could be that RodA3 directly inhibits FtsW1 or displaces FtsW1 at the cell division site, leading to a block in cell division and therefore resulting in the formation of elongated cells.

Recent studies have shown that SEDS proteins act as glycosyltransferases (Meeske et al., 2016, Emami et al., 2017). The glycosyltransferase activity of PBPs and MGT can be inhibited by moenomycin, whereas RodA/FtsW enzymes are not affected by moenomycin and are therefore important for moenomycin resistance (Tamura et al., 1980, Emami et al., 2017). In good agreement with the importance of SEDS proteins for the intrinsic moenomycin resistance, deletion of the genes encoding two of the three RodA enzymes, RodA1 and RodA3, resulted in an increased moenomycin sensitivity of *L. monocytogenes* (Fig. 6C).

In *B. subtilis*, RodA is in a complex with the class B PBP, PBP 2A (also named PbpH), and these two proteins act together to polymerize and crosslink the glycan strands (Wei et al., 2003, Henriques et al., 1998). Similarly, FtsW and PBP 2B form a subcomplex as part of the divisome (Daniel et al., 1996, Gamba et al., 2009). Recently it was shown that RodA-PBP3 and FtsW-PBP1 act as cognate pairs in the coccoid bacterium *S. aureus* (Reichmann et al., 2019). Depletion of all three RodA enzymes in *L. monocytogenes*, RodA1, RodA2 and RodA3, leads to a drastic reduction in cell length (Fig. 5). A similar phenotype was observed for a *L. monocytogenes* strain depleted for the essential class B PBP, PBP B1 (Rismondo et al., 2015). In contrast, the absence of either FtsW1 (Fig. 2) or the class B PBP, PBP B2, in *L. monocytogenes* results in the formation of elongated cells (Rismondo et al., 2015). These observations suggest that RodA and FtsW might work in a complex with the cognate PBPs PBP B1 and PBP B2 during cell elongation and cell division, respectively. Indeed, protein-protein interactions between FtsW1 and FtsW2 with PBP B2 and between RodA1, RodA2 and RodA3 with the PBP B1 could be observed (Fig. 8). However, interactions were also detected between the FtsW proteins and PBP B1 and PBP B3 as well as between the RodA proteins and PBP B2 and PBP B3 (Fig. 8). While these data provide the first line of evidence that SEDS proteins and class B PBPs also form complexes in *L. monocytogenes*, additional work is necessary to determine if specific SEDS-bPBP pairs are formed in *L. monocytogenes*. Taken together, *L. monocytogenes* has a repertoire of PBPs and multiple members of the SEDS family of proteins to produce its rigid cell wall. The expression and the activity of these enzymes need to be tightly regulated in *L. monocytogenes* to maintain its cell shape. Our results suggest that *L. monocytogenes* adapts the expression of a second set of FtsW/RodA enzymes, FtsW2 and RodA3, to environmental stresses such as the presence of β-lactam antibiotics, thereby preventing defects in the peptidoglycan synthesis and subsequent cell lysis. In *L. monocytogenes* strains that lack this back-up system, RodA2 might in part fulfil a similar role.

## Acknowledgements

This work was funded by the Wellcome Trust grants 100289/Z/12/Z and 210671/Z/18/Z to AG and the German research foundation (DFG) grants RI 2920/1-1 to JR and HA6830/1-1 to SH.

---

**Figure S1:**
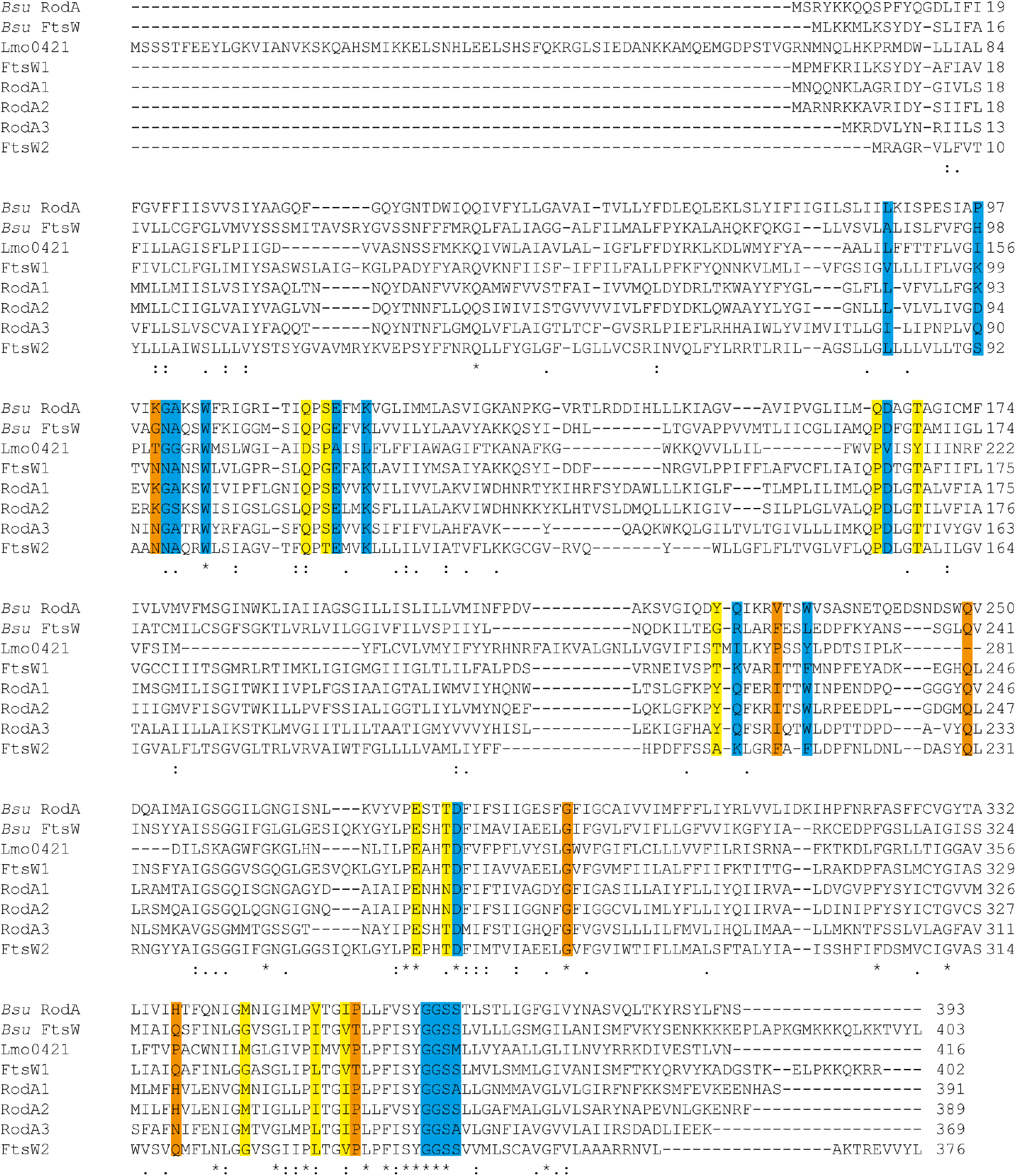
Sequence alignment of FtsW and RodA proteins from *B. subtilis* and *L. monocytogenes*. The alignment was created using Clustal Omega (Sievers et al., 2011). Based on a study by Meeske *et al.*, amino acid residues of *B. subtilis* RodA that are essential for RodA function are colored in blue, where conservative replacements are possible in yellow and amino acids that can be changed to certain other amino acids are colored in orange (Meeske et al., 2016).

**Figure S2:**
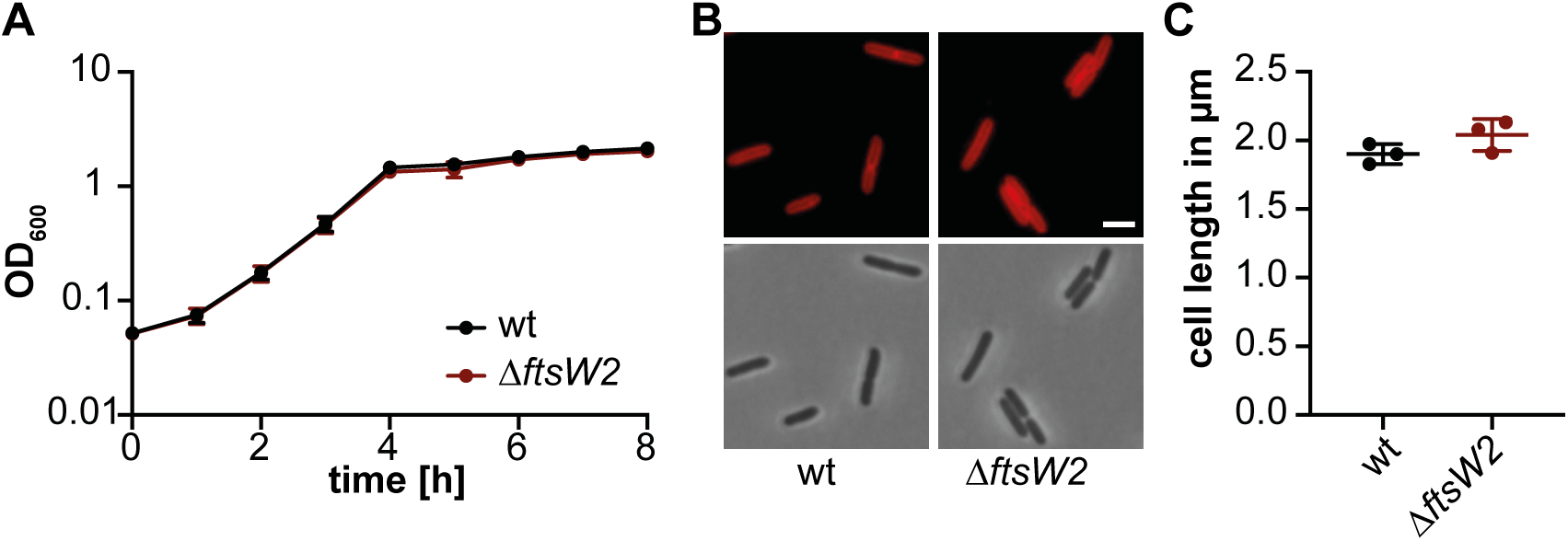
An *L. monocytogenes ftsW2* mutant does not show a growth or cell length defect. (A) Growth of *L. monocytogenes* strains 10403S (wt) and 10403SΔ*ftsW2* in BHI broth at 37°C. (B) Microscopy images of wildtype and *ftsW2* mutant *L. monocytogenes* strains. Bacteria from cultures of the strains described in panel A were stained with the membrane dye nile red and analyzed by fluorescence and phase contrast microscopy. Scale bar is 2 μm. (C) Cell length measurement. The cell length of 300 cells of strain 10403S (wt) and 10403SΔ*ftsW2* was measured and the median cell length calculated and plotted. Three independent experiments were performed, and the average and standard deviation of the median cell length plotted.

**Figure S3:**
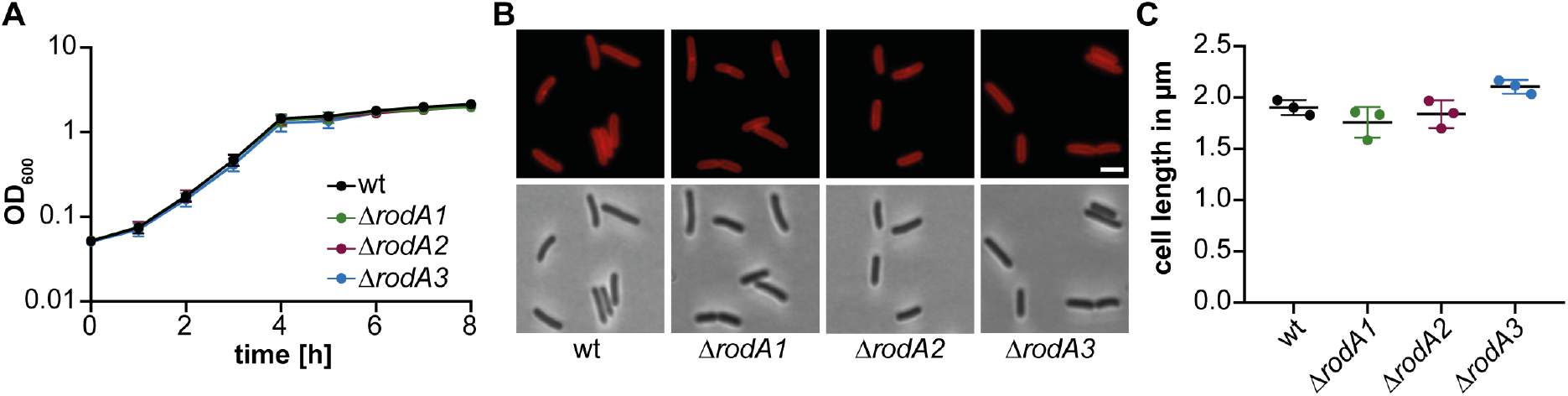
Single *rodA1, rodA2* or *rodA3 L. monocytogenes* strains do not show a growth or cell length defect. (A) Growth of *L. monocytogenes* 10403S (wt), 10403SΔ*rodA1*, 10403SΔ*rodA2* and 10403SΔ*rodA3* in BHI broth at 37°C. (B) Microscopy images of wildtype and mutant *L. monocytogenes* strains. Bacteria from cultures of the strains described in panel A were stained with the membrane dye nile red and analyzed by fluorescence and phase contrast microscopy. Scale bar is 2 μm. (C) Cell length of *L. monocytogenes* wt and *rodA* single mutants. The cell length of 300 cells per strain was measured and the median cell length calculated and plotted. Three independent experiments were performed, and the average and standard deviation of the median cell length plotted.

**Figure S4:**
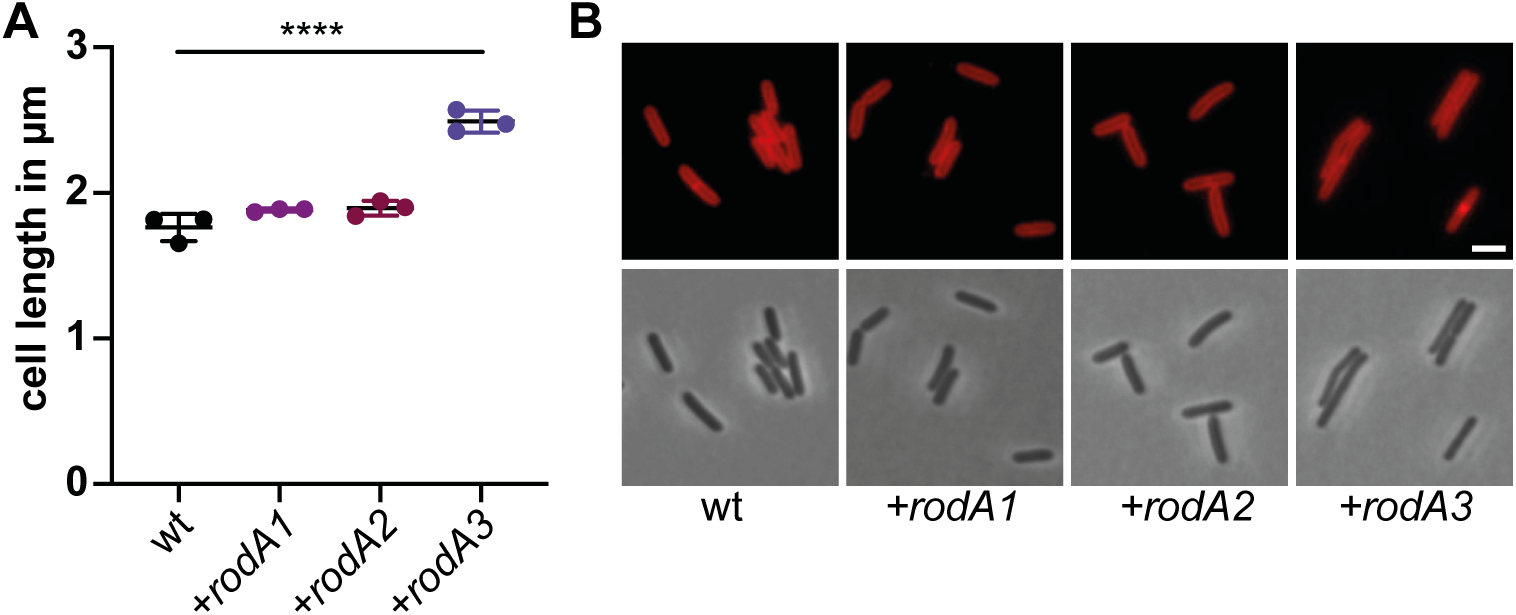
Overexpression of RodA3 results in cell elongation. (A) Cell lengths measurements of 10403S (wt) and strains 10403S pIMK3-*rodA1*, 10403S pIMK3-*rodA2* and 10403S pIMK3-*rodA3*. The cell length of 300 cells per strain was measured. Three independent experiments were performed, and the average and standard deviation of the median cell length plotted. For statistical analysis, a one-way ANOVA coupled with a Dunnett’s multiple comparisons test was performed (**** p≤0.0001). (B) Microscopy images of *L. monocytogenes* strains. Bacteria from cultures of the strains described in panel A were stained with the membrane dye nile red and analyzed by fluorescence and phase contrast microscopy. Scale bar is 2 μm.

**Figure S5:**
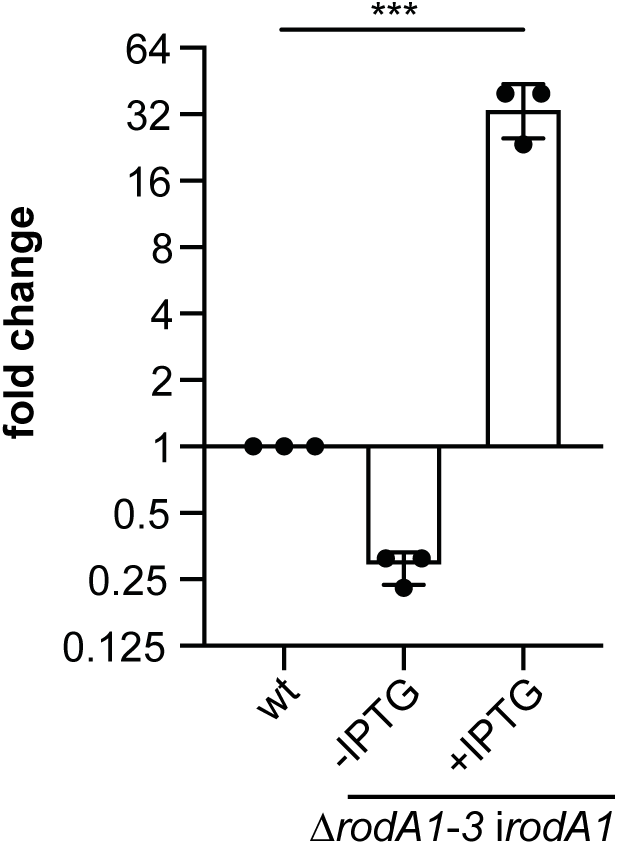
*rodA1* transcripts are still detected even after prolonged RodA1 depletion. Analysis of *rodA1* expression using qRT-PCR. RNA was isolated from strains 10403S and 10403SΔ*rodA1-3* i*rodA1*, that had been grown in absence and presence of IPTG. Expression of *rodA1* was normalized to the expression of *gyrB* and fold changes calculated using the ΔΔCt method. Averages and standard deviations of three independent experiments were plotted. For statistical analysis, a one-way ANOVA coupled with a Dunnett’s multiple comparisons test was performed (*** p≤0.001).

**Figure S6:**
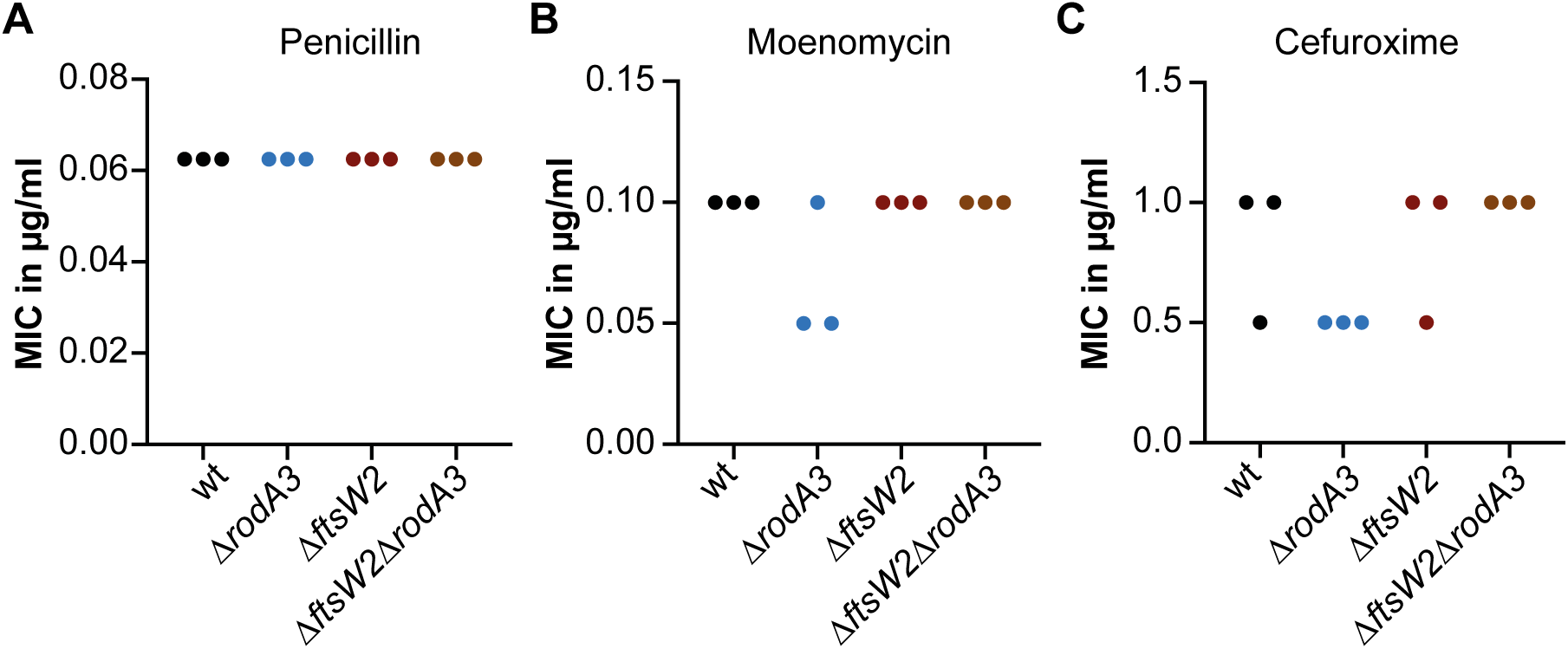
Deletion of *rodA3* but not *ftsW2* has a minor impact on the resistance towards a selected number of cell wall active antibiotics. The minimal inhibitory concentrations for the antibiotics penicillin G (A), moenomycin (B) and cefuroxime (C) were determined for the wildtype *L. monocytogenes* strain 10403S, the 10403SΔ*rodA3* and 10403SΔ*ftsW2* single deletion strains and the 10403SΔ*ftsW2*Δ*rodA3* double mutant strain using a microbroth dilution assay. The result of three biological replicates are shown in A-C.

## References

Aubry, C., Goulard, C., Nahori, M. A., Cayet, N., Decalf, J., Sachse, M., Boneca, I. G., Cossart, P. & Dussurget, O. 2011. OatA, a peptidoglycan O-acetyltransferase involved in *Listeria monocytogenes* immune escape, is critical for virulence. J Infect Dis, 204, 731–40.

Blumberg, P. M. & Strominger, J. L. 1974. Interaction of penicillin with the bacterial cell: penicillin-binding proteins and penicillin-sensitive enzymes. Bacteriol Rev, 38, 291–335.

Boneca, I. G., Dussurget, O., Cabanes, D., Nahori, M. A., Sousa, S., Lecuit, M., Psylinakis, E., Bouriotis, V., Hugot, J. P., Giovannini, M., Coyle, A., Bertin, J., Namane, A., Rousselle, J. C., Cayet, N., Prevost, M. C., Balloy, V., Chignard, M., Philpott, D. J., Cossart, P. & Girardin, S. E. 2007. A critical role for peptidoglycan N-deacetylation in *Listeria* evasion from the host innate immune system. Proc Natl Acad Sci U S A, 104, 997–1002.

Boyle, D. S., Khattar, M. M., Addinall, S. G., Lutkenhaus, J. & Donachie, W. D. 1997. *ftsW* is an essential cell-division gene in *Escherichia coli*. Mol Microbiol, 24, 1263–73.

Burke, T. P., Loukitcheva, A., Zemansky, J., Wheeler, R., Boneca, I. G. & Portnoy, D. A. 2014. *Listeria monocytogenes* is resistant to lysozyme through the regulation, not the acquisition, of cell wall-modifying enzymes. J Bacteriol, 196, 3756–67.

Carballido-Lopez, R. & Formstone, A. 2007. Shape determination in *Bacillus subtilis*. Curr Opin Microbiol, 10, 611–6.

Cho, H., Wivagg, C. N., Kapoor, M., Barry, Z., Rohs, P. D. A., Suh, H., Marto, J. A., Garner, E. C. & Bernhardt, T. G. 2016. Bacterial cell wall biogenesis is mediated by SEDS and PBP polymerase families functioning semi-autonomously. Nature Microbiology, 1, 16172.

Corrigan, R. M., Abbott, J. C., Burhenne, H., Kaever, V. & Gründling, A. 2011. c-di-AMP is a new second messenger in *Staphylococcus aureus* with a role in controlling cell size and envelope stress. PLoS Pathog, 7, e1002217.

Daniel, R. A., Williams, A. M. & Errington, J. 1996. A complex four-gene operon containing essential cell division gene *pbpB* in *Bacillus subtilis*. J Bacteriol, 178, 2343–50.

De Jonge, B. L., Chang, Y. S., Gage, D. & Tomasz, A. 1992. Peptidoglycan composition of a highly methicillin-resistant *Staphylococcus aureus* strain. The role of penicillin binding protein 2A. J Biol Chem, 267, 11248–54.

De Pedro, M. A. & Cava, F. 2015. Structural constraints and dynamics of bacterial cell wall architecture. Front Microbiol, 6, 449.

Den Blaauwen, T. 2018. Is Longitudinal Division in Rod-Shaped Bacteria a Matter of Swapping Axis? Front Microbiol, 9, 822.

Emami, K., Guyet, A., Kawai, Y., Devi, J., Wu, L. J., Allenby, N., Daniel, R. A. & Errington, J. 2017. RodA as the missing glycosyltransferase in *Bacillus subtilis* and antibiotic discovery for the peptidoglycan polymerase pathway. Nat Microbiol, 2, 16253.

Errington, J. & Wu, L. J. 2017. Cell Cycle Machinery in Bacillus subtilis. In: Löwe, J. & Amos, L. A. (eds.) Prokaryotic Cytoskeletons: Filamentous Protein Polymers Active in the Cytoplasm of Bacterial and Archaeal Cells. Cham: Springer International Publishing.

Fraipont, C., Alexeeva, S., Wolf, B., Van Der Ploeg, R., Schloesser, M., Den Blaauwen, T. & Nguyen-Disteche, M. 2011. The integral membrane FtsW protein and peptidoglycan synthase PBP3 form a subcomplex in *Escherichia coli*. Microbiology, 157, 251–9.

Gamba, P., Hamoen, L. W. & Daniel, R. A. 2016. Cooperative Recruitment of FtsW to the Division Site of *Bacillus subtilis*. Front Microbiol, 7, 1808.

Gamba, P., Veening, J. W., Saunders, N. J., Hamoen, L. W. & Daniel, R. A. 2009. Two-step assembly dynamics of the *Bacillus subtilis* divisome. J Bacteriol, 191, 4186–94.

Ghuysen, J. M. & Strominger, J. L. 1963. Structure of the Cell Wall of *Staphylococcus aureus*, Strain Copenhagen. Ii. Separation and Structure of Disaccharides. Biochemistry, 2, 1119–25.

Goffin, C. & Ghuysen, J. M. 1998. Multimodular penicillin-binding proteins: an enigmatic family of orthologs and paralogs. Microbiol Mol Biol Rev, 62, 1079–93.

Gottschalk, S., Bygebjerg-Hove, I., Bonde, M., Nielsen, P. K., Nguyen, T. H., Gravesen, A. & Kallipolitis, B. H. 2008. The two-component system CesRK controls the transcriptional induction of cell envelope-related genes in *Listeria monocytogenes* in response to cell wall-acting antibiotics. J Bacteriol, 190, 4772–6.

Gründling, A., Burrack, L. S., Bouwer, H. G. & Higgins, D. E. 2004. *Listeria monocytogenes* regulates flagellar motility gene expression through MogR, a transcriptional repressor required for virulence. Proc Natl Acad Sci U S A, 101, 1231823.

Hara, H. & Suzuki, H. 1984. A novel glycan polymerase that synthesizes uncross-linked peptidoglycan in *Escherichia coli*. FEBS Lett, 168, 155–60.

Henriques, A. O., De Lencastre, H. & Piggot, P. J. 1992. A *Bacillus subtilis* morphogene cluster that includes *spoVE* is homologous to the *mra* region of *Escherichia coli*. Biochimie, 74, 735–48.

Henriques, A. O., Glaser, P., Piggot, P. J. & Moran, C. P., Jr. 1998. Control of cell shape and elongation by the *rodA* gene in *Bacillus subtilis*. Mol Microbiol, 28, 235–47.

Höltje, J. V. 1998. Growth of the stress-bearing and shape-maintaining murein sacculus of *Escherichia coli*. Microbiol Mol Biol Rev, 62, 181–203.

Ikeda, M., Sato, T., Wachi, M., Jung, H. K., Ishino, F., Kobayashi, Y. & Matsuhashi, M. 1989. Structural similarity among *Escherichia coli* FtsW and RodA proteins and *Bacillus subtilis* SpoVE protein, which function in cell division, cell elongation, and spore formation, respectively. J Bacteriol, 171, 6375–8.

Jones, L. J., Carballido-Lopez, R. & Errington, J. 2001. Control of cell shape in bacteria: helical, actin-like filaments in *Bacillus subtilis*. Cell, 104, 913–22.

Kallipolitis, B. H., Ingmer, H., Gahan, C. G., Hill, C. & Sogaard-Andersen, L. 2003. Cesrk, a two-component signal transduction system in *Listeria monocytogenes*, responds to the presence of cell wall-acting antibiotics and affects beta-lactam resistance. Antimicrob Agents Chemother, 47, 3421–9.

Karimova, G., Pidoux, J., Ullmann, A. & Ladant, D. 1998. A bacterial two-hybrid system based on a reconstituted signal transduction pathway. Proc Natl Acad Sci U S A, 95, 5752–6.

Karinou, E., Schuster, C. F., Pazos, M., Vollmer, W. & Gründling, A. 2018. Inactivation of the monofunctional peptidoglycan glycosyltransferase SgtB allows *Staphylococcus aureus* to survive in the absence of lipoteichoic acid. J Bacteriol.

Khattar, M. M., Begg, K. J. & Donachie, W. D. 1994. Identification of FtsW and characterization of a new *ftsW* division mutant of *Escherichia coli*. J Bacteriol, 176, 7140–7.

Kobayashi, K., Ehrlich, S. D., Albertini, A., Amati, G., Andersen, K. K., Arnaud, M., Asai, K., Ashikaga, S., Aymerich, S., Bessieres, P., Boland, F., Brignell, S. C., Bron, S., Bunai, K., Chapuis, J., Christiansen, L. C., Danchin, A., Debarbouille, M., Dervyn, E., Deuerling, E., Devine, K., Devine, S. K., Dreesen, O., Errington, J., Fillinger, S., Foster, S. J., Fujita, Y., Galizzi, A., Gardan, R., Eschevins, C., Fukushima, T., Haga, K., Harwood, C. R., Hecker, M., Hosoya, D., Hullo, M. F., Kakeshita, H., Karamata, D., Kasahara, Y., Kawamura, F., Koga, K., Koski, P., Kuwana, R., Imamura, D., Ishimaru, M., Ishikawa, S., Ishio, I., Le Coq, D., Masson, A., Mauel, C., Meima, R., Mellado, R. P., Moir, A., Moriya, S., Nagakawa, E., Nanamiya, H., Nakai, S., Nygaard, P., Ogura, M., Ohanan, T., O’Reilly, M., O’Rourke, M., Pragai, Z., Pooley, H. M., Rapoport, G., Rawlins, J. P., Rivas, L. A., Rivolta, C., Sadaie, A., Sadaie, Y., Sarvas, M., Sato, T., Saxild, H. H., Scanlan, E., Schumann, W., Seegers, J. F., Sekiguchi, J., Sekowska, A., Seror, S. J., Simon, M., Stragier, P., Studer, R., Takamatsu, H., Tanaka, T., Takeuchi, M., Thomaides, H. B., Vagner, V., Van Dijl, J. M., Watabe, K., Wipat, A., Yamamoto, H., Yamamoto, M., Yamamoto, Y., Yamane, K., Yata, K., Yoshida, K., Yoshikawa, H., Zuber, U. & Ogasawara, N. 2003. Essential *Bacillus subtilis* genes. Proc Natl Acad Sci U S A, 100, 4678–83.

Korsak, D., Markiewicz, Z., Gutkind, G. O. & Ayala, J. A. 2010. Identification of the full set of *Listeria monocytogenes* penicillin-binding proteins and characterization of PBPD2 (Lmo2812). BMC Microbiol, 10, 239.

Leclercq, S., Derouaux, A., Olatunji, S., Fraipont, C., Egan, A. J., Vollmer, W., Breukink, E. & Terrak, M. 2017. Interplay between Penicillin-binding proteins and SEDS proteins promotes bacterial cell wall synthesis. Sci Rep, 7, 43306.

Liu, X., Gallay, C., Kjos, M., Domenech, A., Slager, J., Van Kessel, S. P., Knoops, K., Sorg, R. A., Zhang, J. R. & Veening, J. W. 2017. High-throughput CRISPRi phenotyping identifies new essential genes in *Streptococcus pneumoniae*. Mol Syst Biol, 13, 931.

Lobel, L. & Herskovits, A. A. 2016. Systems Level Analyses Reveal Multiple Regulatory Activities of CodY Controlling Metabolism, Motility and Virulence in *Listeria monocytogenes*. PLoS Genet, 12, e1005870.

Mcpherson, D. C. & Popham, D. L. 2003. Peptidoglycan Synthesis in the absence of class A penicillin-binding proteins in *Bacillus subtilis*. J Bacteriol, 185, 1423–31.

Meeske, A. J., Riley, E. P., Robins, W. P., Uehara, T., Mekalanos, J. J., Kahne, D., Walker, S., Kruse, A. C., Bernhardt, T. G. & Rudner, D. Z. 2016. SEDS proteins are a widespread family of bacterial cell wall polymerases. Nature, 537, 634–638.

Meeske, A. J., Sham, L. T., Kimsey, H., Koo, B. M., Gross, C. A., Bernhardt, T. G. & Rudner, D. Z. 2015. MurJ and a novel lipid II flippase are required for cell wall biogenesis in *Bacillus subtilis*. Proc Natl Acad Sci U S A, 112, 6437–42.

Nakagawa, J., Tamaki, S. & Matsuhashi, M. 1979. Purified penicillin binding proteins 1Bs from *Escherichia coli* membrane showing activities of both peptidoglycan polymerase and peptidoglycan crosslinking enzyme. Agric. Biol. Chem., 43, 1379–1380.

Nanninga, N. 1991. Cell division and peptidoglycan assembly in *Escherichia coli*. Mol Microbiol, 5, 791–5.

Nielsen, P. K., Andersen, A. Z., Mols, M., Van Der Veen, S., Abee, T. & Kallipolitis, B. H. 2012. Genome-wide transcriptional profiling of the cell envelope stress response and the role of LisRK and CesRK in *Listeria monocytogenes*. Microbiology, 158, 963–74.

Park, W. & Matsuhashi, M. 1984. *Staphylococcus aureus* and *Micrococcus luteus* peptidoglycan transglycosylases that are not penicillin-binding proteins. J Bacteriol, 157, 538–44.

Park, W., Seto, H., Hakenbeck, R. & Matsuhashi, M. 1985. Major peptidoglycan transglycosylase activity in *Streptococcus pneumoniae* that is not a penicillin-binding protein. FEMS Microbiology Letters, 27, 45–48.

Pinho, M. G., Kjos, M. & Veening, J. W. 2013. How to get (a)round: mechanisms controlling growth and division of coccoid bacteria. Nat Rev Microbiol, 11, 601–14.

Reichmann, N. T., Tavares, A. C., Saraiva, B. M., Jousselin, A., Reed, P., Pereira, A. R., Monteiro, J. M., Sobral, R. G., Vannieuwenhze, M. S., Fernandes, F. & Pinho, M. G. 2019. SEDS-bPBP pairs direct lateral and septal peptidoglycan synthesis in *Staphylococcus aureus*. Nat Microbiol.

Rismondo, J., Möller, L., Aldridge, C., Gray, J., Vollmer, W. & Halbedel, S. 2015. Discrete and overlapping functions of peptidoglycan synthases in growth, cell division and virulence of *Listeria monocytogenes*. Mol Microbiol, 95, 332–51.

Rogers, H. J., Perkins, H. R. & Ward, J. B. 1980. Microbial Cell Walls and Membranes. Chapman and Hall New York, N.Y., 10, 33–33.

Ruiz, N. 2008. Bioinformatics identification of MurJ (MviN) as the peptidoglycan lipid II flippase in *Escherichia coli*. Proc Natl Acad Sci U S A, 105, 15553–7.

Sauvage, E., Kerff, F., Terrak, M., Ayala, J. A. & Charlier, P. 2008. The Penicillin-binding proteins: structure and role in peptidoglycan biosynthesis. FEMS Microbiol Rev, 32, 234–58.

Scheffers, D. J. & Pinho, M. G. 2005. Bacterial cell wall synthesis: new insights from localization studies. Microbiol Mol Biol Rev, 69, 585–607.

Sham, L. T., Butler, E. K., Lebar, M. D., Kahne, D., Bernhardt, T. G. & Ruiz, N. 2014. Bacterial cell wall. MurJ is the flippase of lipid-linked precursors for peptidoglycan biogenesis. Science, 345, 220–2.

Sievers, F., Wilm, A., Dineen, D., Gibson, T. J., Karplus, K., Li, W., Lopez, R., Mcwilliam, H., Remmert, M., Soding, J., Thompson, J. D. & Higgins, D. G. 2011. Fast, scalable generation of high-quality protein multiple sequence alignments using Clustal Omega. MolSyst Biol, 7, 539.

Sonnhammer, E. L., Von Heijne, G. & Krogh, A. 1998. A hidden Markov model for predicting transmembrane helices in protein sequences. Proc Int Conf Intell Syst Mol Biol, 6, 175–82.

Stone, K. J. & Strominger, J. L. 1971. Mechanism of action of bacitracin: complexation with metal ion and C 55-isoprenyl pyrophosphate. Proc Natl Acad Sci U S A, 68, 3223–7.

Swaminathan, B. & Gerner-Smidt, P. 2007. The epidemiology of human listeriosis. Microbes Infect, 9, 1236–43.

Taguchi, A., Welsh, M. A., Marmont, L. S., Lee, W., Sjodt, M., Kruse, A. C., Kahne, D., Bernhardt, T. G. & Walker, S. 2019. FtsW is a peptidoglycan polymerase that is functional only in complex with its cognate penicillin-binding protein. Nat Microbiol, 4, 587–594.

Tamura, T., Suzuki, H., Nishimura, Y., Mizoguchi, J. & Hirota, Y. 1980. On the process of cellular division in *Escherichia coli*: isolation and characterization of penicillin-binding proteins 1a, 1b, and 3. Proc Natl Acad Sci U S A, 77, 4499–503.

Toledo-Arana, A., Dussurget, O., Nikitas, G., Sesto, N., Guet-Revillet, H., Balestrino, D., Loh, E., Gripenland, J., Tiensuu, T., Vaitkevicius, K., Barthelemy, M., Vergassola, M., Nahori, M. A., Soubigou, G., Regnault, B., Coppee, J. Y., Lecuit, M., Johansson, J. & Cossart, P. 2009. The *Listeria* transcriptional landscape from saprophytism to virulence. Nature, 459, 950–6.

Typas, A., Banzhaf, M., Gross, C. A. & Vollmer, W. 2012. From the regulation of peptidoglycan synthesis to bacterial growth and morphology. Nat Rev Microbiol, 10, 123–36.

Van Heijenoort, J. 2001. Formation of the glycan chains in the synthesis of bacterial peptidoglycan. Glycobiology, 11, 25R–36R.

Van Heijenoort, Y., Leduc, M., Singer, H. & Van Heijenoort, J. 1987. Effects of moenomycin on *Escherichia coli*. J Gen Microbiol, 133, 667–74.

Vollmer, W., Blanot, D. & De Pedro, M. A. 2008. Peptidoglycan structure and architecture. FEMS Microbiol Rev, 32, 149–67.

Wang, Q. M., Peery, R. B., Johnson, R. B., Alborn, W. E., Yeh, W. K. & Skatrud, P. L. 2001. Identification and characterization of a monofunctional glycosyltransferase from *Staphylococcus aureus*. J Bacteriol, 183, 4779–85.

Wei, Y., Havasy, T., Mcpherson, D. C. & Popham, D. L. 2003. Rod shape determination by the *Bacillus subtilis* class B penicillin-binding proteins encoded by *pbpA* and *pbpH*. J Bacteriol, 185, 4717–26.

Weidel, W. & Pelzer, H. 1964. Bagshaped Macromolecules--a New Outlook on Bacterial Cell Walls. Adv Enzymol Relat Areas Mol Biol, 26, 193–232.

Zhang, C., Nietfeldt, J., Zhang, M. & Benson, A. K. 2005. Functional consequences of genome evolution in *Listeria monocytogenes*: the *lmo0423* and *lmo0422* genes encode sigmaC and LstR, a lineage II-specific heat shock system. J Bacteriol, 187, 7243–53.

