## Supplementary Methods for "Cell shape and antibiotic resistance is maintained by the activity of multiple FtsW and RodA enzymes in *Listeria monocytogenes*"

### Strain and plasmid construction

All strains and primers used in this study are listed in Tables S1 and S2, respectively. For the construction of markerless in-frame deletions of the *L. monocytogenes* genes *ftsW1* (*lmo1071*), *rodA2* (*lmo2428*), and *ftsW2* (*lmo2688*), 1-kb DNA fragments of the up- and downstream regions of the respective genes were amplified by PCR with primer pairs ANG2440/2441 and ANG2442/2443 (*ftsW1*), ANG2456/2457 and 2458/2459 (*rodA2*) or ANG2444/2445 and ANG2446/2447 (*ftsW2*). The resulting PCR products were fused using primers ANG2440/2443 (*ftsW1*), ANG2456/2459 (*rodA2*) and ANG2444/2447 (*ftsW2*), cut with BamHI and KpnI and ligated with the plasmid pKSV7 that had been cut with the same enzymes. The resulting plasmids pKSV7-Δ*rodA2* and pKSV7-Δ*ftsW2* were recovered in *E. coli* XL1-Blue yielding strains ANG4132 and ANG4137, respectively. pKSV7-Δ*ftsW1* was recovered in *E. coli* CLG190 yielding strain ANG4169. For the generation of pKSV7-Δ*rodA1* and pKSV7-Δ*rodA3*, up- and downstream fragments of *rodA1* and *rodA3* were amplified using primers ANG2452/2453 and ANG2454/2455 (*rodA1*) or ANG2448/2832 and ANG2833/2451 (*rodA3*) and used as the template for a second PCR using primers ANG2452/2455 (*rodA1*) and ANG2448/2451 (*rodA3*), respectively. The resulting PCR fragments were digested with XbaI and KpnI and ligated with the plasmid pKSV7 that had been cut with the same enzymes. The resulting plasmids pKSV7-Δ*rodA1* and pKSV7-Δ*rodA3* were recovered in *E. coli* XL1-Blue yielding strains ANG4135 and ANG4665, respectively. For the deletion of the two-gene operons *rodA1-2* and *ftsW2-rodA3,* 1kb-DNA fragments of the up- and downstream region of the operon were amplified using primer pairs ANG2721/2722 and ANG2723/2455 (*rodA1-2*) or ANG2444/2719 and ANG2720/2742 (*ftsW2-rodA3*). The resulting PCR products were fused using primers ANG2721/2455 (*rodA1-2*) and ANG2444/2742 (*ftsW2-rodA3*), cut with BamHI and KpnI and ligated with pKSV7 also cut with BamHI/KpnI. Plasmids pKSV7-Δ*rodA1*Δ*rodA2* and pKSV7-Δ*ftsW2*Δ*rodA3* were recovered in *E. coli* XL1-Blue yielding strains ANG4444 and ANG4457, respectively. The pKSV7-derivatives were subsequently electroporated into the *L. monocytogenes* strain 10403S and the genes deleted by allelic exchange as described previously (Camilli et al., 1993). The deletion of the respective genes was verified by PCR resulting in the construction of strains 10403SΔ*rodA1* (ANG4171), 10403SΔ*rodA2* (ANG4172), 10403SΔ*rodA3* (ANG4683), 10403SΔ*ftsW2* (ANG4176), 10403SΔ*rodA1*Δ*rodA2* (ANG4459) and 10403SΔ*ftsW2*Δ*rodA3* (ANG4482). Subsequently, *rodA3* was deleted in strains 10403SΔ*rodA1* and 10403SΔ*rodA2* yielding strains 10403SΔ*rodA1*Δ*rodA3* (ANG4684) and 10403SΔ*rodA2*Δ*rodA3* (ANG4685). For complementation analysis of strain 10403SΔ*rodA1*Δ*rodA3*, plasmids pIMK3-*rodA1*, pIMK3-*rodA2* and pIMK3-*rodA3* were constructed allowing for IPTG-dependent gene expression. For this purpose, *rodA1, rodA2* and *rodA3* were amplified using primers ANG2951/2952, ANG3008/3009 and ANG2953/2954, respectively. The *rodA1* and *rodA3* PCR products were digested with NcoI and BamHI and the *rodA2* PCR product was digested with NcoI and SalI. The digested PCR products were subsequently ligated with plasmid pIMK3 that had been cut with the corresponding enzymes. The resulting plasmids were recovered in XL1-Blue yielding strains ANG4892 (pIMK3-*rodA1*), ANG4976 (pIMK3-*rodA2*) and ANG4893 (pIMK3-*rodA3*). Next, plasmids pIMK3-*rodA1,* pIMK3-*rodA2* or pIMK3-*rodA3* were introduced into *L. monocytogenes* strain 10403SΔ*rodA1*Δ*rodA3* yielding strains 10403SΔ*rodA1*Δ*rodA3* pIMK3-*rodA1* (ANG4905), 10403SΔ*rodA1*Δ*rodA3* pIMK3-*rodA2* (ANG5149) and

10403SΔ*rodA1*Δ*rodA3* pIMK3-*rodA3* (ANG4906). In addition, plasmids pIMK3-*rodA1,* pIMK3-*rodA2* or pIMK3-*rodA3* were transformed into *L. monocytogenes* strain 10403S yielding strains 10403S pIMK3-*rodA1* (ANG4966), 10403S pIMK3-*rodA2* (ANG4981) and 10403S pIMK3-*rodA3* (ANG4967).

Despite undertaking several attempts, we were unable to construct a strain in which all three *rodA* genes were deleted in *L. monocytogenes* 10403S, indicating that at least one *rodA* gene needs to be present. Therefore, we generated a strain in which *rodA1* is expressed from an ectopic, IPTG-inducible promoter and deleted all three *rodA* genes from the genome. For this purpose, plasmid pIMK3-*rodA1* (from strain ANG4892) was transformed into *L. monocytogenes* 10403SΔ*rodA1*Δ*rodA2* (ANG4459) yielding strain 10403SΔ*rodA1*Δ*rodA2* pIMK3-*rodA1* (ANG5148). Subsequently, *rodA3* was deleted from the genome of strain 10403SΔ*rodA1*Δ*rodA2* pIMK3-*rodA1* (ANG5148) by allelic exchange in the presence of 1 mM IPTG using plasmid pKSV7-Δ*rodA3.* The resulting strain 10403SΔ*rodA1-3* pIMK3-*rodA1* (ANG5192) was verified by PCR.

Attempts to delete *ftsW1* in *L. monocytogenes* 10403S remained unsuccessful, indicating that *ftsW1* is an essential gene. Therefore, a plasmid to facilitate the ectopic, IPTG-inducible expression of *ftsW1* was constructed. For this purpose, the *ftsW1* gene was amplified using primers ANG2594/2595, the product cut with NcoI and SalI and ligated with plasmid pIMK3 that had been cut with the same enzymes. The resulting plasmid was recovered in *E. coli* XL1-Blue yielding strain ANG4261. pIMK3-*ftsW1* was subsequently electroporated into *L. monocytogenes* 10403S yielding strain 10403S pIMK3-*ftsW1* (ANG4288). For the construction of a conditional *ftsW1* deletion strain, plasmid pKSV7-Δ*ftsW1* was introduced into strain 10403S pIMK3-*ftsW1* (ANG4288) and the allelic exchange procedure was carried out in the presence of 1 mM IPTG. The resulting strain 10403SΔ*ftsW1* pIMK3-*ftsW1* (or 10403S i*ftsW1*, ANG4314) was verified by PCR. To determine whether FtsW2 can take over the role of FtsW1, we constructed a strain in which the expression of *ftsW2* can be induced by IPTG. For this purpose, *ftsW2* was amplified using primers ANG3010/3011, the product cut with NcoI and SalI and ligated with plasmid pIMK3 that had been cut with the same enzymes. Plasmid pIMK3-*ftsW2* was recovered in XL1-Blue yielding strain ANG4977. The plasmid was subsequently introduced into *L. monocytogenes* 10403S yielding strain 10403S pIMK3-*ftsW2* (ANG4982). In the next step, pKSV7-Δ*ftsW1* was introduced into strain ANG4982 and the allelic exchange procedure was carried out in the presence of IPTG. The resulting strain 10403SΔ*ftsW1* pIMK3-*ftsW2* (or 10403SΔ*ftsW1* i*ftsW2*; ANG5119) was verified by PCR.

For the construction of plasmid pPL3e-P$_{lmo2689}$-*lacZ*, the promoterless *lacZ* gene was amplified from plasmid p*itet*-P$_{700\text{-}ltaS}$-*lacZ* (ANG2027) using primers ANG3131/3132, digested with SalI/KpnI and ligated with plasmid pPL3e that had been cut with the same enzymes. The resulting plasmid pPL3e-*lacZ* was recovered in *E. coli* XL1-Blue yielding strain ANG5181. In the next step, primers ANG3141 and ANG3142 were used to amplify an approximately 600 bp region upstream of *lmo2689*, including the promoter region and the first 9 bases of *lmo2689*. The resulting PCR product was cut with BamHI/SalI and ligated with plasmid pPL3e-*lacZ*. Plasmid pPL3e-P$_{lmo2689}$-*lacZ* was recovered in *E. coli* XL1-Blue yielding strain ANG5193 and was subsequently introduced by electroporation into *L. monocytogenes* strains 10403S, 10403SΔ*rodA1* and 10403SΔ*rodA1*Δ*rodA2* yielding strains ANG5198, ANG5292 and ANG5293, respectively.

For bacterial two-hybrid assays, *ftsW1, ftsW2, rodA1, rodA2* and *rodA3* were amplified using primer pairs JR341/342, JR343/344, JR347/348, JR345/346 and JR349/350, respectively. The resulting PCR products were cut with XbaI and KpnI and ligated with plasmids pUT18 and pUT18c that had been cut with the same enzymes. The resulting plasmids pUT18-*ftsW1* (pJR206), pUT18c-*ftsW1* (pJR212), pUT18-*ftsW2* (pJR205), pUT18c-*ftsW2* (pJR211), pUT18-*rodA1* (pJR201), pUT18c-*rodA1* (pJR207), pUT18-*rodA2* (pJR204), pUT18c-*rodA2* (pJR210), pUT18-*rodA3* (pJR202) and pUT18c-*rodA3* (pJR208) were recovered in *E. coli* TOP10.

CAMILLI, A., TILNEY, L. G. & PORTNOY, D. A. 1993. Dual roles of *plcA* in *Listeria monocytogenes* pathogenesis. *Mol Microbiol,* 8**,** 143-57.
