## Supplementary Table S1 for "Cell shape and antibiotic resistance is maintained by the activity of multiple FtsW and RodA enzymes in *Listeria monocytogenes*"

**Table S1: Bacterial strains used in this study**

| Unique ID | Strain name and resistance | Source |
|---|---|---|
| ***Escherichia coli* strains** | | |
| ANG1264 | DH5α pKSV7; AmpR | (Smith and Youngman, 1992) |
| ANG1278 | XL1-Blue pPL3e; CamR | (Gründling et al., 2004) |
| ANG2027 | XL1-Blue p*itet*-$P_{700\text{-}ltaS}$-*lacZ*, AmpR | Lab strain collection |
| ANG4243 | XL1-Blue pIMK3; KanR | (Monk et al., 2008) |
| ANG4132 | XL1-Blue pKSV7-Δ*rodA2*; AmpR | This study |
| ANG4135 | XL1-Blue pKSV7-Δ*rodA1*; AmpR | This study |
| ANG4137 | XL1-Blue pKSV7-Δ*ftsW2*; AmpR | This study |
| ANG4169 | CLG190 pKSV7-Δ*ftsW1*; AmpR | This study |
| ANG4261 | XL1-Blue pIMK3-*ftsW1*; KanR | This study |
| ANG4444 | XL1-Blue pKSV7-Δ*rodA1*Δ*rodA2*; AmpR | This study |
| ANG4457 | XL1-Blue pKSV7-Δ*ftsW2*Δ*rodA3*; AmpR | This study |
| ANG4665 | XL1-Blue pKSV7-Δ*rodA3*; AmpR | This study |
| ANG4892 | XL1-Blue pIMK3-*rodA1*; KanR | This study |
| ANG4893 | XL1-Blue pIMK3-*rodA3*; KanR | This study |
| ANG4976 | XL1-Blue pIMK3-*rodA2*; KanR | This study |
| ANG4977 | XL1-Blue pIMK3-*ftsW2*; KanR | This study |
| ANG5181 | XL1-Blue pPL3e-*lacZ*; CamR | This study |
| ANG5193 | XL1-Blue pPL3e-$P_{600\text{-}lmo2689}$-*lacZ*; CamR | This study |
| ANG5402 | XL1 Blue pKT25; KanR | (Karimova et al., 1998) |
| ANG5415 | XL1 Blue pUT18; AmpR | (Karimova et al., 1998) |
| ANG5416 | XL1 Blue pUT18c; AmpR | (Karimova et al., 1998) |
| pJR201 | TOP10 pUT18-*rodA1*; AmpR | This study |
| pJR202 | TOP10 pUT18-*rodA3*; AmpR | This study |
| pJR204 | TOP10 pUT18-*rodA2*; AmpR | This study |
| pJR205 | TOP10 pUT18-*ftsW2*; AmpR | This study |
| pJR206 | TOP10 pUT18-*ftsW1*; AmpR | This study |
| pJR207 | TOP10 pUT18c-*rodA1*; AmpR | This study |
| pJR208 | TOP10 pUT18c-*rodA3*; AmpR | This study |
| pJR210 | TOP10 pUT18c-*rodA2*; AmpR | This study |
| pJR211 | TOP10 pUT18c-*ftsW2*; AmpR | This study |
| pJR212 | TOP10 pUT18c-*ftsW1*; AmpR | This study |
| pSH235 | TOP10 pKT25-*pbpB2*; KanR | (Cleverley et al., 2019) |
| pSH236 | TOP10 pKT25-*pbpB1*; KanR | (Cleverley et al., 2019) |
| pSH237 | TOP10 pKT25-*pbpB3*; KanR | (Cleverley et al., 2019) |

| | | |
|---|---|---|
| ***Listeria monocytogenes* strains** | | |
| ANG1263 | 10403S; StrepR | (Bishop and Hinrichs, 1987) |
| ANG4171 | 10403SΔ*rodA1*; StrepR | This study |
| ANG4172 | 10403SΔ*rodA2*; StrepR | This study |
| ANG4176 | 10403SΔ*ftsW2*; StrepR | This study |
| ANG4288 | 10403S pIMK3-*ftsW1*; StrepR KanR | This study |
| ANG4314 | 10403SΔ*ftsW1* pIMK3-*ftsW1*; StrepR KanR, IPTG | This study |
| ANG4459 | 10403SΔ*rodA1*Δ*rodA2*; StrepR | This study |
| ANG4482 | 10403SΔ*ftsW2*Δ*rodA3*; StrepR | This study |
| ANG4683 | 10403SΔ*rodA3*; StrepR | This study |
| ANG4684 | 10403SΔ*rodA1*Δ*rodA3*; StrepR | This study |
| ANG4685 | 10403SΔ*rodA2*Δ*rodA3*; StrepR | This study |
| ANG4905 | 10403SΔ*rodA1*Δ*rodA3* pIMK3-*rodA1*; StrepR KanR | This study |
| ANG4906 | 10403SΔ*rodA1*Δ*rodA3* pIMK3-*rodA3*; StrepR KanR | This study |
| ANG4966 | 10403S pIMK3-*rodA1*; StrepR KanR | This study |
| ANG4967 | 10403S pIMK3-*rodA3*; StrepR KanR | This study |
| ANG4981 | 10403S pIMK3-*rodA2*; StrepR KanR | This study |
| ANG4982 | 10403S pIMK3-*ftsW2*; StrepR KanR | This study |
| ANG5119 | 10403SΔ*ftsW1* pIMK3-*ftsW2*; StrepR KanR, IPTG | This study |
| ANG5148 | 10403SΔ*rodA1*Δ*rodA2* pIMK3-*rodA1*; StrepR KanR | This study |
| ANG5149 | 10403SΔ*rodA1*Δ*rodA3* pIMK3-*rodA2*; StrepR KanR | This study |
| ANG5192 | 10403SΔ*rodA1-3* pIMK3-*rodA1*; StrepR KanR, IPTG | This study |
| ANG5198 | 10403S pPL3e-$P_{lmo2689}$-*lacZ*; StrepR ErmR | This study |
| ANG5292 | 10403SΔ*rodA1* pPL3e-$P_{lmo2689}$-*lacZ*; StrepR ErmR | This study |
| ANG5293 | 10403SΔ*rodA1*Δ*rodA2* pPL3e-$P_{lmo2689}$-*lacZ*; StrepR ErmR | This study |


BISHOP, D. K. & HINRICHS, D. J. 1987. Adoptive transfer of immunity to *Listeria monocytogenes*. The influence of in vitro stimulation on lymphocyte subset requirements. *J Immunol,* 139**,** 2005-9.

CLEVERLEY, R. M., RUTTER, Z. J., RISMONDO, J., CORONA, F., TSUI, H. T., ALATAWI, F. A., DANIEL, R. A., HALBEDEL, S., MASSIDDA, O., WINKLER, M. E. & LEWIS, R. J. 2019. The cell cycle regulator GpsB functions as cytosolic adaptor for multiple cell wall enzymes. *Nat Commun,* 10**,** 261.

GRÜNDLING, A., BURRACK, L. S., BOUWER, H. G. & HIGGINS, D. E. 2004. *Listeria monocytogenes* regulates flagellar motility gene expression through MogR, a transcriptional repressor required for virulence. *Proc Natl Acad Sci U S A,* 101**,** 12318-23.

KARIMOVA, G., PIDOUX, J., ULLMANN, A. & LADANT, D. 1998. A bacterial two-hybrid system based on a reconstituted signal transduction pathway. *Proc Natl Acad Sci U S A,* 95**,** 5752-6.

MONK, I. R., GAHAN, C. G. & HILL, C. 2008. Tools for functional postgenomic analysis of *Listeria monocytogenes*. *Appl Environ Microbiol,* 74**,** 3921-34.

SMITH, K. & YOUNGMAN, P. 1992. Use of a new integrational vector to investigate compartment-specific expression of the *Bacillus subtilis spoIIM* gene. *Biochimie,* 74**,** 705-11.
