## Supplementary Table S2 for "Cell shape and antibiotic resistance is maintained by the activity of multiple FtsW and RodA enzymes in *Listeria monocytogenes*"

**Table S2: Primers used in this study**

| Number | Name | Sequence |
|---|---|---|
| ANG2440 | Lmo1071_up_fw | GCGCGGATCCGCGTACCAGATCCAAGTAGATTTG |
| ANG2441 | Lmo1071_up_rev | CTTTGGTAACTCATCATAAGATTTTAAAATTCGTTTAAACAT |
| ANG2442 | Lmo1071_down_fw | ATCTTATGATGAGTTACCAAAGAAACAAAAAAGACGTTAA |
| ANG2443 | Lmo1071_down_rev | GCGCGGTACCGCAGCATCACAGATTCGGTTACGTAG |
| ANG2444 | Lmo2688_up_fw | GCGCGGATCCCATCGACGAAAGAAGCAATTGCTCATC |
| ANG2445 | Lmo2688_up_rev | TCGTTTTAGCCGTTACAAATAGTACGCGTCCAGCACGCAT |
| ANG2446 | Lmo2688_down_fw | ATTTGTAACGGCTAAAACGAGAGAGGTCGTGTATTTATGA |
| ANG2447 | Lmo2688_down_rev | GCGCGGTACCGAATGGAAGCGGAATCCCCGTGAG |
| ANG2448 | Lmo2687_up_fw | GCGCTCTAGAGGTGTTGCTGTTATGCGATATAAAGTG |
| ANG2451 | Lmo2687_down_rev | GCGCGGTACCGTTAGAAGTCATTTCCGTTGTGAGAGTTC |
| ANG2452 | Lmo2427_up_fw | GCGCTCTAGAGATAAGCTGCAATGGGCGGCATAC |
| ANG2453 | Lmo2427_up_rev | CTTTCACTTCGCGTCCAGCTAGTTTATTTTGTTGATTCAT |
| ANG2454 | Lmo2427_down_fw | AGCTGGACGCGAAGTGAAAGAAGAAAATCACGCATCTTAA |
| ANG2455 | Lmo2427_down_rev | GCGCGGTACCCCCAAGCCGGCCTCCTAAAAATTAATC |
| ANG2456 | Lmo2428_up_fw | GCGCGGATCCCCAGTTGGAATCTTGCTTGCAATG |
| ANG2457 | Lmo2428_up_rev | CTAAGTTAACTCGAACTGCCTTTTTTCTATTTCTGGCCAT |
| ANG2458 | Lmo2428_down_fw | GGCAGTTCGAGTTAACTTAGGGAAAGAAAATCGTTTCTGA |
| ANG2459 | Lmo2428_down_rev | GCGCGGTACCCGGTACGCCAACATCCAGCGCCAC |
| ANG2594 | Lmo1071_fw_NcoI | GCGCCCATGGGGCCGATGTTTAAACGAATTTTAA |
| ANG2595 | Lmo1071_rev_SalI | GCGCGTCGACTTAACGTCTTTTTTGTTTCTTTGGTA |
| ANG2719 | Lmo2688/7 up rev | ATCGGCATCTGACGTTACAAATAGTACGCGTCCAGCACGCAT |
| ANG2720 | Lmo2688/7 down fw | CTATTTGTAACGTCAGATGCCGATTTAATAGAAGAAAAATAA |
| ANG2721 | Lmo2428/7 up fw | GCGCGGATCCCCAGTTGGAATCCTTCTTGCAATG |
| ANG2722 | Lmo2428/7 up rev | TTCTTTCACTTCTCGAACTGCCTTTTTTCTATTTCTGGCCAT |
| ANG2723 | Lmo2428/7 down fw | AAGGCAGTTCGAGAAGTGAAAGAAGAAAATCACGCATCTTAA |
| ANG2742 | Lmo2688/7 down rev | GCGCGGTACCGATACCAATAACTAAAACAGCTTC |

| | | |
|---|---|---|
| ANG2832 | Lmo2687 up rev new | ATCGGCATCTGAAATTCGGTTATAAAGTACATCTCGTTTCAT |
| ANG2833 | Lmo2687 down fw new | TATAACCGAATTTCAGATGCCGATTTAATAGAAGAAAAATAA |
| ANG2951 | pIMK3-2427 fw NcoI | GCGCCCATGGGGAATCAACAAAATAAACTAGCTGG |
| ANG2952 | pIMK3-2427 rev Bam | GCGCGGATCCTTAAGATGCGTGATTTTCTTC |
| ANG2953 | pIMK3-2687 fw NcoI | GCGCCCATGGGGAAACGAGATGTACTTTATAACC |
| ANG2954 | pIMK3-2687 rev Bam | GCGCGGATCCTTATTTTTCTTCTATTAAATCGGC |
| ANG3008 | pIMK3 2428 fw Nco | GCGCCCATGGCCAGAAATAGAAAAAAGGCAG |
| ANG3009 | pIMK3 2428 rev Sal | GCGCGTCGACTCAGAAACGATTTTCTTTCCCTAAG |
| ANG3010 | pIMK3 2688 NcoI | GCGCCCATGGGGCGTGCTGGACGCGTACTA |
| ANG3011 | pIMK3 2688 SalI | GCGCGTCGACTCATAAATACACGACCTCTCTC |
| ANG3131 | pPL3e lacZ fw SalI | ACGCGTCGACCGGGAAAACCCTG |
| ANG3132 | pPL3e lacZ rev KpnI | CGGGGTACCTTATTTTTGACACCAGACCAACTG |
| ANG3141 | P600-2689 fw | GCGCGGATCCCAACCGCAGATTCATTAGCAG |
| ANG3142 | P600-2689 rev | GCGCGTCGACTAGTTTTTTCATTTTCCTTGCCTCC |
| JR341 | B2H lmo1071 fw | GCGCTCTAGAATTTAAACGAATTTTAAAATCTTATGATTATGC |
| JR342 | B2H lmo1071 rev | GCGCGGTACCGCACGTCTTTTTTGTTTCTTTGGTAACTC |
| JR343 | B2H lmo2688 fw | GCGCTCTAGAACGTGCTGGACGCGTACTATTTG |
| JR344 | B2H lmo2688 rev | GCGCGGTACCGCTAAATACACGACCTCTCTCGTTTTAG |
| JR347 | B2H lmo2427 fw | GCGCTCTAGAAAATCAACAAAATAAACTAGCTGGACG |
| JR348 | B2H lmo2427 rev | GCGCGGTACCGCAGATGCGTGATTTTCTTCTTTCACTTC |
| JR349 | B2H lmo2687 fw | GCGCTCTAGAAAAACGAGATGTACTTTATAACCGG |
| JR350 | B2H lmo2687 rev | GCGCGGTACCGCTTTTTCTTCTATTAAATCGGCATCTG |
